# Integrative multi-omics analysis reveals molecular subtypes and tumor evolution of synovial sarcoma

**DOI:** 10.1101/2022.05.09.490894

**Authors:** Yi Chen, Yanhong Su, Isabelle Rose Leo, Ioannis Siavelis, Jianming Zeng, Xiaofang Cao, Panagiotis Tsagkozis, Asle C Hesla, Andri Papakonstantinou, Xiao Liu, Wen-Kuan Huang, Monika Ehnman, Henrik Johansson, Yingbo Lin, Janne Lehtiö, Yifan Zhang, Olle Larsson, Felix Haglund de Flon

## Abstract

Synovial sarcomas (SS) are malignant mesenchymal tumors characterized by the SS18-SSX fusion gene, which drives tumorigenesis by altering the composition of the BAF complex. Secondary genomic alterations that determine variations in tumor phenotype or clinical presentation are largely unknown. Herein, we present transcriptome, targeted DNA-sequencing, and proteomics analysis of 91 synovial sarcomas from 55 patients. We identified three SS clusters (SSCs) characterized by distinct histology, tumor microenvironments, genomic complexities, therapeutic effects, and clinical outcomes. Eight BAF complex components are differentially expressed among SSCs, and their role in mesenchymal-epithelial-transition is supported by single cell sequencing. The epithelial cells of biphasic tumors are more susceptible to developing copy number alterations, including amplification of *PDCD1* and *TMPRSS2*. Our findings explain broad concepts in SS biology and imply that the BAF composition at the start of the tumorigenesis (i.e. the cellular linage) may determine the SS subtype, providing a rationale for individualized treatment strategies.

## Introduction

Synovial sarcoma (SS) is an aggressive mesenchymal soft tissue malignancy. It represents the fourth most common type of soft tissue sarcoma (STS), accounting for 5–10% of all primary STS worldwide[1–4]. All SSs are characterized by a chromosomal translocation t(X;18)(p11;q11). This rearrangement results in a fusion between the SS18 gene on chromosome 18 (encoding a member of the chromatin modeling BAF complex) and one of three homologous genes *SSX1*, *SSX2,* or *SSX4* on chromosome X that form the fusion protein *SS18-SSX* [5–8]. The chimeric oncoprotein *SS18-SSX* hijacks the BAF complex and consequently reprograms chromatin architecture to promote sarcomagenesis [9–14], essentially gaining stem-cell-like properties. Histologically SS can appear as monophasic (purely mesenchymal) or biphasic (mesenchymal and epithelial). These two subtypes are almost equally distributed (55% vs. 45%, respectively) [15, 16]. Cases with increased nuclear atypia, necrosis, and higher mitotic index can be classified as poorly differentiated variants [15, 16]. However, apart from studying the primary fusion gene SS18-SSX, the limited number of comprehensively studied cases constrains the understanding of the secondary genomic events in SS[17–19].

SS is a relatively chemosensitive subtype compared to other STS, but the efficacy of conventional therapeutic agents seems limited in patients with metastatic disease [20, 21]. A substantial proportion of patients with localized tumors treated with curative resection, adjuvant chemotherapy, and radiotherapy will develop either local relapse or distant metastasis within a short time period [22–24]. On the other hand, other patients with only surgical treatment may never relapse. In addition, SS is considered to be a poorly immunogenic type of cancer, and immune checkpoint inhibitors (ICIs) have thus far shown a low response rate in the treatment of SS [25, 26]. The BAF complex is a major chromatin regulator which can mediate resistance to ICIs in cancers [27, 28]. The unfavorable therapeutic effectiveness in the metastatic setting strongly motivates new therapies and a thorough understanding of the SS landscape at the transcriptome, genome, and proteome levels.

We hypothesized that the unknown spectrum of secondary genetic events could help distinguish relevant biological subgroups of SS to aid clinicians in determining the prognosis and optimal treatment of individual patients. Thus, we set out to comprehensively characterize SS tumors from a multi-omics perspective treated at our institution for the last 30 years.

## Results

### Clinical and tumor characteristics of SS patients

A schematic presentation of the study design is represented in Fig. 1a. Among the 55 SS patients, 88 tumor samples had both whole transcriptomic data (WTS) and target gene DNA-seq data (Extended Data Fig. 1a-b), of which 14 fresh frozen samples had proteomics data. As shown in Supplementary Table 1, seven of 21 patients (33%) that received neoadjuvant therapy had >10% viable tumor tissue (poor responder). A total of 28 (50.9%) patients developed distant metastases (median: 5.1 years, range: 0 – 24 years), of which 24 (43.6%) patients died of the disease during the follow-up. Patients with metastasis were given chemotherapy and radiotherapy (16.7%), chemotherapy alone (29.2%), or radiotherapy alone (37.5%). The 5-year overall survival rate was 50.9%, and the 10-year overall survival rate was 41.5%, confirming the aggressive nature of SS.

**Figure 1:**
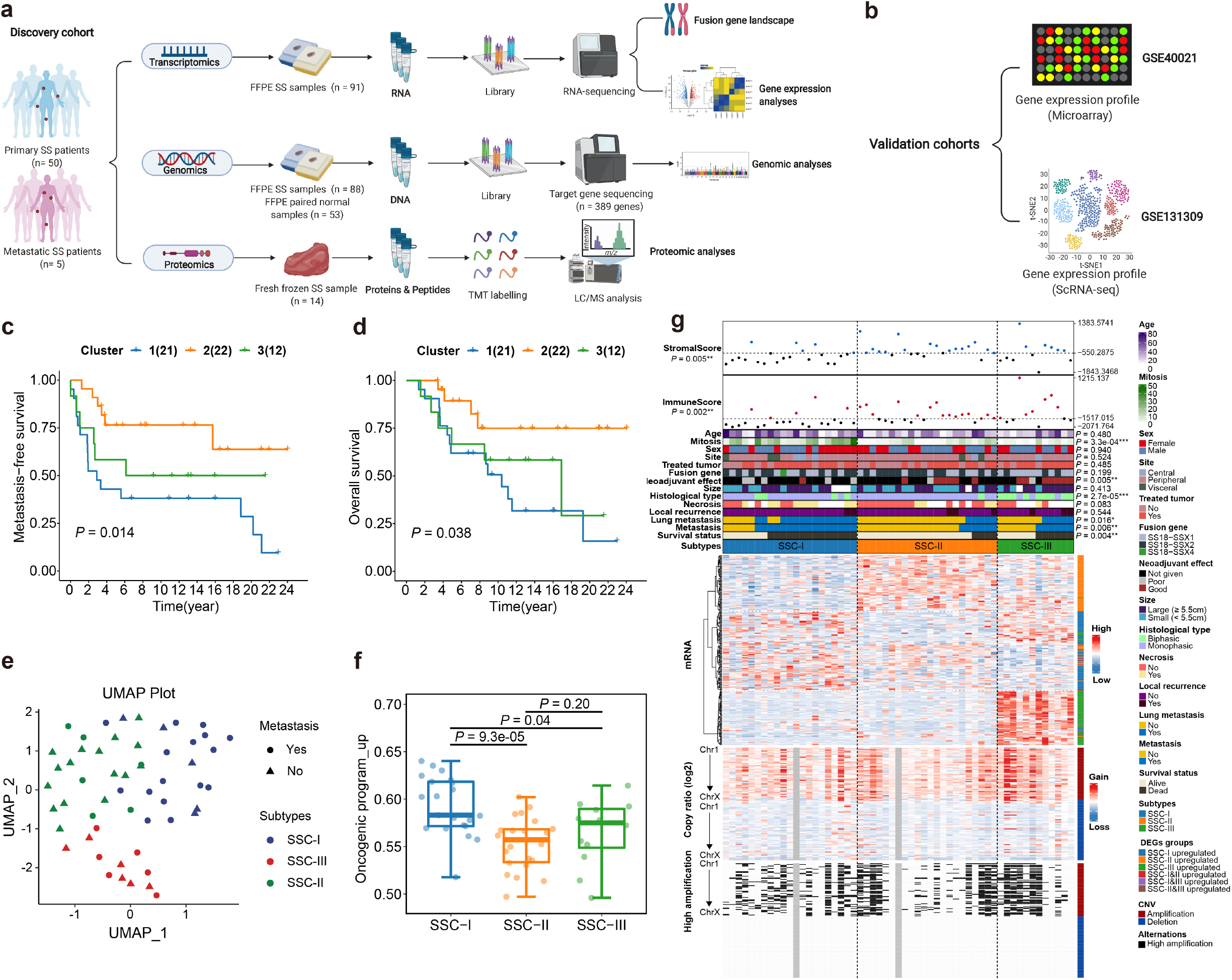
Synovial sarcoma subtypes. **(a-b).** The study workflow, including **(a)** discovery and **(b)** validation cohorts. **(c).** Kaplan-Meier curve of clusters of metastasis-free survival in synovial sarcoma patients with cluster outliers incorporated at top 14% gene variance (Log-rank test, *P* = 0.014). **(d).** Kaplan-Meier curve of overall survival in synovial sarcoma patients with cluster outliers incorporated at top 14% gene variance (Log-rank test, *P* = 0.038). **(e).** UMAP plot of SS subtypes and metastatic status distributions. **(f).** Distributions of the ssGSEA score of upregulated genes in the oncogenic program across three SSCs. Middle line: median; box edges: 25th and 75th percentiles. Mann-Whitney U test. **(G).** Heatmap of the three SSCs. Clinicopathological characteristics (top) of the 55 synovial sarcoma patients are shown in the annotation, and different colors represent the characteristics and subtypes. Molecular data (bottom) from 55 patients profiled with mRNA and copy number variations are depicted. The statistical differences in categorical variables with three subtypes were compared using Fisher’s exact test; Continuous variables were compared using the Kruskal-Wallis test. \**p* < 0.05, \*\**p* < 0.01, \*\*\**p* < 0.001.

### Fusion gene identification and the SS18-SSX fusion gene landscape in SS

Gene fusions were detected in all 91 RNA-sequenced samples from 55 patients by the FusionCatcher (primary) and STAR-Fusion (supportive) pipelines. Prior to filtering, there were 9750 candidate fusion transcripts with the FusionCatcher tool across all the samples, with *SS18-SSX* detected in all but three samples. The SS18-SSX chimeric gene was confirmed in the RNA-sequenced tissue using the chimeric-protein-specific antibody (Extended Data Fig. 1c).

Since previous studies have shown that variants of the *SS18-SSX1*, *SS18-SSX2*, and *SS18-SSX4* were observed in SS [29], we sought to characterize the *SS18-SSX* fusion variants in our cohort. The usual fusions were present in most of the cases, whereas a novel *SS18-SSX2* fusion variant was detected involving the traditional breakpoint in *SS18* transcripts (…ATATGACCAG*) and novel *SSX2* (*GACCCAAAG…) on ENST0000033677 (Extended Data Fig. 2a). As expected, the *SS18-SSX1* and *SS18-SSX2* variants predominated, and the *SS18-SSX4* occurred in only seven samples (Extended Data Fig. 2a). Next, we wanted to assess the expression level of the fusion gene in ten patients with multiple samples in a longitudinal analysis. By assessing the raw expression of *SSX1-4,* we were able to conclude that the transcripts of the SSX genes originated from the fusion gene and not the normal allele. Expression levels of *SS18* and *SSX* (corresponding to the correct fusion gene) were plotted and compared to the clinical course and treatments. We found an inconspicuous change in *SS18* expression after radiotherapy or chemotherapy, while the expression level of *SSX1* dropped in most cases after treatment or metastasis, but this was less noticeable in the case of *SSX2* (Extended Data Fig. 2b). Recent studies suggested that the fusion variant does not hold prognostic value [30, 31]. The clinical relevance of the involved SSX gene was investigated by Kaplan-Meier plots, and our data also supported that there was no significant difference between SSX1 and SSX2 fusions in terms of overall survival (OS) or metastasis-free survival (MFS) (log-rank test, *P* = 0.637 and 0.494, respectively) (Extended Data Fig. 2c), similar to the patient outcomes in the validation cohort (log-rank test, *P* = 0.498, Extended Data Fig. 2d).

Given the high noise due to FFPE samples and the low specificity of the fusion gene identification pipelines, a rigorous filtering process was performed in the FusionCatcher (Extended Data Fig. 3 and Extended Data Fig. 4a), excluding banned (n = 979), high common mapping reads (n = 1108), promiscuous genes (n = 5459), low bioinformatics support (n = 1529), adjacent (n = 23), and short repeats (n = 110), which were considered to be false positives and were removed before downstream analyses. After filtering and IGV-inspection, Fusion Catcher identified 115 putative true fusion genes, including 96 in-frame fusions, eight fusions with reciprocal reads, nine with supporting exonic reads, and two previously identified fusions (Mitelman/ChimerDB databases). STAR-Fusion appeared to be less sensitive than FusionCatcher in our benchmark analysis, as the SS18-SSX fusion genes were detected only in 74 (81%) of our samples. After a series of filtering processes to exclude adjacent genes (n = 108), very local rearrangements (n = 41), non-reference splice sites (n = 267), “promiscuous” genes (n = 1), and procadherins (n = 2), a total of 29 fusion genes were classified as putative true events (Extended Data Fig. 3 and Extended Data Fig. 4b). There were only five overlapping fusion events between the STAR-Fusion and FusionCatcher whitelists (Extended Data Fig. 3).

To explore the relevance of secondary fusion genes, the whitelisted fusion genes were projected to further validation at the molecular level. No secondary fusion genes were associated with the SS18 or SSX chromosomal regions (Extended Data Fig. 3, Supplementary table 2). We found that 41 fusion genes arose from intrachromosomal rearrangements, while 98 of 139 fusion genes were potentially generated by interchromosomal translocations. Notably, in-frame fusions comprised most fusion pairs, making it easy to exert biologically functional protein production. We also investigated if the number of secondary fusion genes was associated with treatment or metastatic disease, but no significant associations could be found (Extended Data Fig. 4d-e). RT-PCR successfully validated 12 fusion genes (Extended Data Fig. 4f), six of which were successfully cloned and sequenced, including two BAZ2B-DNAH11 variants (Extended Data Fig. 4g). Interestingly, the BAZ2B-DNAH11, EDA-KDM5C, NOR1-SPAG9, and ZFHX4-LRRC69 fusion genes had protein functions associated with chromatin remodeling. However, none of the whitelisted fusion genes were recurrent in our cohort (Extended Data Fig. 5a-f), suggesting that secondary fusion events are rare, perhaps random, and with limited clinical importance. These findings support previous data showing that the primary fusion gene SS18-SSX is the main driver of tumorigenesis in SS.

### Molecular classification and transcriptomic characteristics of SS

To unveil the tumor heterogeneity of SS, we performed non-negative matrix factorization (NMF) approach based on the transcriptomic data of mRNAs in 50 primary and five metastatic patients. A total of 25 thresholds were set (from top 6% - top 30% variant mRNAs) (Extended Data Fig. 6a). According to the optimal K numbers in each group, patients were classified into given groups (Extended Data Fig. 6b-c). Noticeably, all 25 comparisons had outliers (groups with ≤ 3 patients). Three different MFS rates were computed, including original, outliers incorporated, and outliers removed (see Methods for more details). Based on the *p*-value (log-rank test, *p* < 0.05) compared in all three groups, three candidates were retained to observe their clustering consensus matrix and overall survival rate (top 14%, top 16%, and top 23% variance) (Extended Data Fig. 6d). As a result, compared to the top 16% variance and top 23% variance (Extended Data Fig. 7a and 7d), the consensus matrix showed that the top 14% had less overlap between clusters (Extended Data Fig. 7g). The silhouette plots also indicated good clustering of samples and genes in the top 14% variance group (Extended Data Fig. 7i). Additionally, after incorporating the outliers, no significant difference was seen in overall survival for both the top 16% (Extended Data Fig. 7c) and the top 23% (Extended Data Fig. 7f) variance groups (log-rank test, *P* = 0.058 and 0.082, respectively), regardless of the significance of MFSs (log-rank test, *P* < 0.05) (Extended Data Fig. 7b and 7e). Nonetheless, in the top 14% variance group, prognosis was significantly different regarding both MFSs and OS (log-rank test, *P* < 0.05) (Extended Data Fig. 7h; Fig. 1c-d). To further assess the performance of the three models, receiver operating characteristic (ROC) analysis was performed. As shown in Extended Data Fig. 7j, the top 14% variance model had the best areas under the curve (AUC) in both MFS and OS (AUC = 0.733 and 0.735, respectively). Thus, it was selected as the most robust consensus NMF clustering for the downstream analysis.

To further explore the effectiveness and robustness of our three SS clusters (SSC-I, II, and III), we visualized metastatic and primary samples using Uniform Manifold Approximation and Projection (UMAP), which showed a clear separation of the three groups except for the metastasis group (Fig. 1e). We then examined the top 14% variant genes in the validation cohort (GSE40021), which had consistent characteristics where cluster 1 (blue) denotes the worst prognosis, and cluster 2 (orange) represents the low-risk group (log-rank test, *P* = 0.001) (Extended Data Fig. 7k-m). Components of the SS *core oncogenic program* [22] associated with poor prognosis, anti-immunity, and aggressive disease were found to be frequently amplified in multiple SS cohorts [32]. To investigate this *core oncogenic program* in the three identified SS subtypes, *core oncogenic program* genes enrichment scores were calculated by Single-sample Gene Set Enrichment Analysis (ssGSEA) methodology using GSVA [33]. Interestingly, upregulation of the core oncogenic program was significantly higher in SSC-I, followed by SSC-III and SSC-II (Mann-Whitney test, *P* = 0.04 and *P* = 9.3e-05, respectively) (Fig. 1f). Likewise, in the GEO cohort, cluster1 was mostly enriched in the upregulated program, and least in the downregulated program (Mann-Whitney test, *P* < 0.05, respectively) (Extended Data Fig. 7n-o). Subsequently, we investigated the four significant proliferative hallmark gene sets between SSC-I, II, and III. As expected, SSC-I was mostly enriched in proliferative gene sets, and SSC-III had the lowest score (Extended Data Fig. 7p). When comparing the ssGSEA scores in immune pathways, SSC-I showed the lowest immune presence, while SSC-III exhibited the highest immune response (Extended Data Fig. 7q).

Next, we applied multiple immune infiltration computing algorithms to access the tumor microenvironment in SS. In the pan-cancer analysis based on the ESTIMATE method, SS showed an intermediate abundance of stromal components (*stromal score*) but exceptionally low immunological component (*immune score*) (Extended Data Fig. 8a). In terms of specific immune cell proportions based on the CIBERSORT algorithms, particularly high levels of naïve B cells and CD4 T cells in SS and low infiltration of memory B cells and T cells, indicating immune-evasion mechanisms in line with previous studies [22, 34] (Extended Data Fig. 8b). When comparing differences between metastatic and primary SS for immune- and stromal cells, few were significant, (Extended Data Fig. 9a-c), perhaps indicative of immune-independent proliferative pathways. During the follow-up period (range 16-292 months), 24 patients died, all of which had distant metastasis, predominately lung metastasis, and to a lesser extent lymph node metastasis and rarely other organs. To identify gene expression patterns associated with metastatic disease, we performed five comparisons (see Methods) for GSEA. All comparisons showed the high enrichment score in gene sets associated with proliferation (E2F targets, cell cycle, mitosis, mitochondrial translation, etc., normalized *P* < 0.001, FDR < 0.1), which is in line with the SSC-I transcriptomic patterns (Supplementary Table 6). Enriched gene sets overlapping the comparisons were identified and included four hallmark gene sets, seven genes ontology-biological process (GO-BP), and 20 Reactome pathways (Extended Data Fig. 9d-g), suggesting that proliferation is a key mechanism associated with metastasis.

### Functional annotations of three SSCs

Considering the substantial differences seen between the three subtypes, we sought to compare the clinical and transcriptomic characteristics of the groups and determine the similarities and differences, as well as to explore the strengths of our classification system and its capability to reveal SS heterogeneity (Fig. 1g). We observed that SSC-I was significantly associated with poor survival (Fisher’s exact test, *P* = 0.004), metastasis (Fisher’s exact test, *P* = 0.006), had significantly highest rates of mitosis (Kruskal-Wallis test, *P* = 3.3e-04) and the lowest stromal and immune scores (calculated by the ESTIMATE algorithm) (Kruskal-Wallis test, *P* = 0.005 and 0.002 respectively). SSC-II, the least aggressive group, had the lowest frequency of death and metastasis and was histologically dominated by monophasic cases (Fisher’s exact test, *P* = 2.7e-05) with the highest stromal score. On the contrary, biphasic cases dominated SSC-III, which also had the highest estimated immune score. Notably, many patients in SSC-III had good response to neoadjuvant treatment (Fisher’s exact test, *P* = 0.005), however most of them later metastasized. There was no significant association with age (Kruskal-Wallis test, *P* > 0.05), sex, site, fusion gene type, size or local recurrence (Fisher’s exact test, *P* > 0.05). The top 50 DEGs (logFC >1.5, adjust *p*.value < 0.01) between each group were assembled, and we performed functional enrichment analysis for the DEGs generated by six comparisons (Extended Data Fig. 9h-j). SSC-I was characterized by the hyperproliferative features and SSC-II with vascular-related pathways. The DEGs between SSC-III and SSC-I were related to innate immune response. SSC-III had a higher expression of cell junction and epithelial cell differentiations compared to SSC-II, which reflected the different histological patterns. At the genomic level, SSC-III showed the highest level of copy number alterations, which may explain the different transcriptome and phenotype of SSC-III.

To further investigate the co-expression profiles of the subtypes, samples were clustered and gene co-expression modules were identified with a soft threshold of 5 for the scale-free network using the weighted gene co-expression network analysis (WGCNA) (Extended Data Fig. 10a). As a result, a total of 15 gene modules were generated according to the dissimilarity measure (1-TOM), where the gray modules are collections of genes that cannot be aggregated to other modules (Extended Data Fig. 10b-c). Furthermore, the relationships between each module and subtypes were evaluated using Spearman’s rank correlation. As shown in Fig. 2a, the purple module revealed a significant positive correlation with SSC-I (ρ = 0.43, *P* =0.001) and a negative correlation with SSC-II (ρ = −0.47, *P* = 3e-04). On the other hand, magenta showed the opposite result (ρ = −0.63, *P* = 2e-07, in SSC-I; *r* = 0.69, *P* = 5e-09, in SSC-II). Remarkably, in SSC-III, the blue module showed the strongest positive correlation (ρ = 0.88, *P* = 2e-18), but negative correlations in SSC-I (ρ = −0.39, *P* = 0.003), and II (ρ = −0.35, *P* = 0.009). The brown (ρ = 0.46, *P* = 4e-04) and black (ρ = 0.47, *P* = 3e-04) modules were highly correlated with SSC-III, but negatively correlated with SSC-I (ρ = −0.55, *P* = 1e-05) and SSC-II (ρ = −0.67, *P* = 2e-05), respectively. The correlation between the gene significance and the module membership score was all significant in the five modules (ρ = 0.37, *P* = 3e-05 in purple; ρ = 0.67, *P* = 3e-23 in magenta; ρ = 0.68, *P* = 3.5e-26 in black; ρ = 0.77, *P* = 1.3e-135 in blue; ρ = 0.43, *P* = 3.6e-28 in brown), illustrating that genes significantly associated with our subtypes correspond to the most important elements of these modules (Extended Data Fig. 10d).

**Figure 2:**
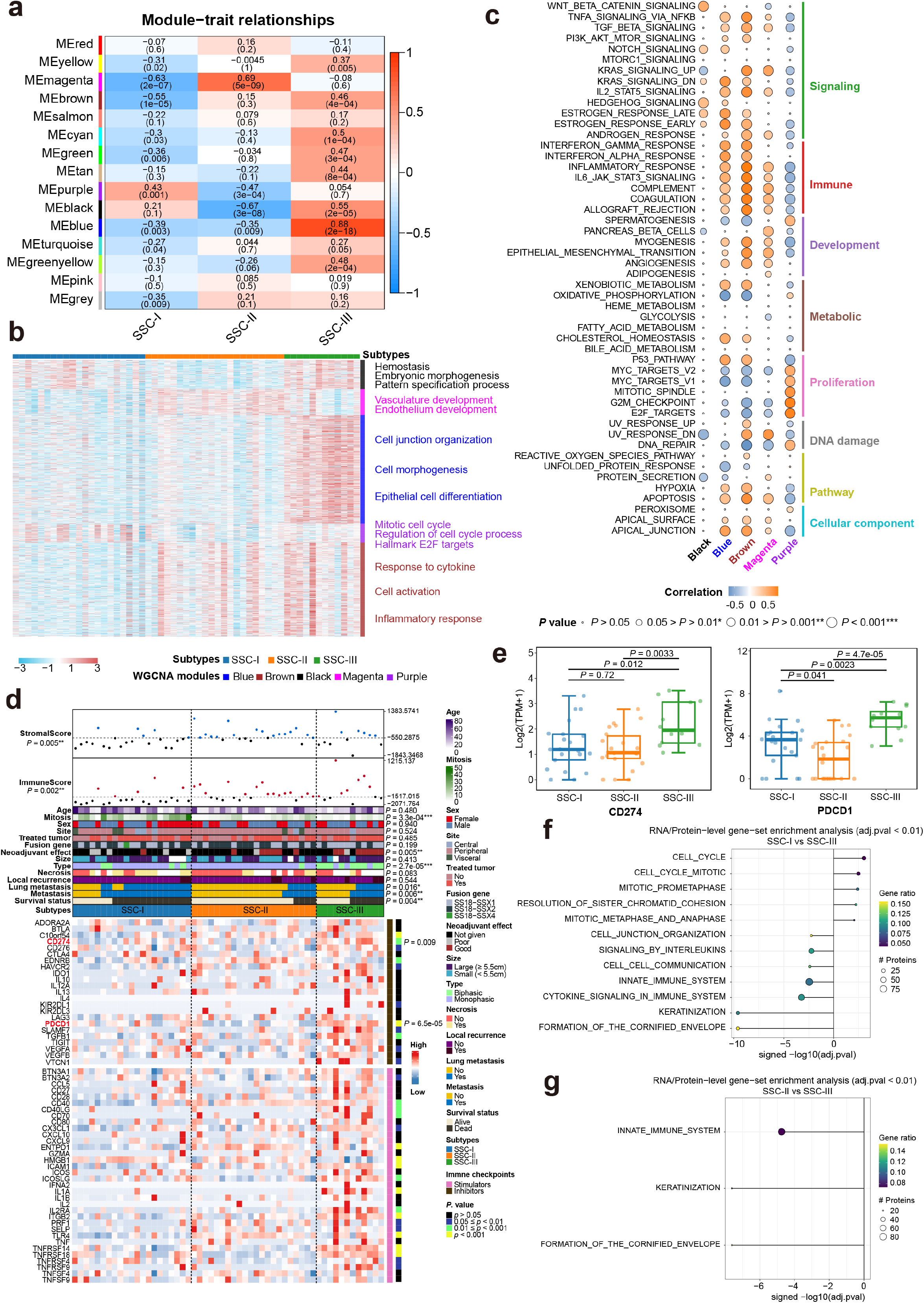
Functional annotation of three SSCs. **(A).** Relationships between WCGNA gene modules and three SSCs. Spearman rank correlation. **(B).** The abundance of five gene modules (black, magenta, blue, purple, and brown) in three SSCs and their top functional annotations on the right. **(C**) Association of five gene modules with 50 Hallmarks on ssGSEA scores. Spearman rank correlation. The shade of the color represents the levels of correlations, the size of circles represents the *p-*value. **(d).** Heatmap of the three SSCs on the mRNA levels of 23 inhibitory and 36 stimulatory immune checkpoints. Clinicopathological characteristics (top) of the 55 SS patients are shown in the annotation, and different colors represent the characteristics and subtypes. The statistical differences in categorical variables were compared using the Fisher’s exact test; Continues variables were compared using Kruskal-Wallis test. \**p* < 0.05, \*\**p* < 0.01, \*\*\**p* < 0.001. **(e)** Distributions of the gene expressions of *CD274* and *PDCD1* in three synovial sarcoma subtypes. Middle line: median; box edges: 25th and 75th percentiles. Mann-Whitney U test. **(f-g).** The functional enrichment between **(f)** SSC-I, **(g)** SSC-II and SSC-III in the RNA/protein level. Gene ratio: In the enrichment analysis, the number of gene symbols (overlapping RNAs and proteins) at the leading edge is divided by the total number of gene symbols in the given pathway. The size of circles represents the number of enriched proteins at the leading edge.

Next, we examined the functional enrichment among differently regulated genes in identified modules. As expected, many significantly enriched genes were associated with specific biological functions, including cell cycle (purple), vasculature and endothelium development (magenta), cell junction and epithelial cell differentiation (blue), immune response (brown), and the functional enrichment analysis of the genes within each module revealed striking concordances with the DEGs analysis (Fig. 2b, Extended Data Fig. 9h-j and 10e). To comprehensively uncover the biological roles of these key modules, the ssGSEA approach was applied for these modules in each patient and detected the correlations with 50 hallmarks. As shown in Fig. 2c, the purple module showed association of high proliferative pathways as well as DNA damage, but resistance to immunity and signaling, indicating proliferative and malignant features in SSC-I, which might be reflective of a high level of treatment resistance (poor effect of neoadjuvant therapy) (Fig. 1g). For magenta, we noticed that the elevated correlation with down-regulated genes in response to ultraviolet (UV) radiation, and low DNA repair capacity, might suggest an SSC-II phenotype of with high drug sensitivity and limited aggressiveness. More complex associations could be observed between the SSC-III phenotype and three representative gene modules. There was distinct functional relevance between black and blue/brown modules. Multiple apoptosis and immune response pathways were significantly upregulated in SSC-III, including IFN-alpha and IFN-gamma, suggesting a high innate immunity response. In the black module, we observed associations to Wnt/β-catenin signaling, KRAS-signaling downregulation, and UV response.

By implementing three deconvolutional algorithms and cross-validation, we profiled the immune landscape across three subtypes. The highest stromal score and immune score were shown in SSC-II and SSC-III, respectively (Extended Data Fig. 11a). The proportion of endothelial cells was highest in SSC-II (Kruskal-Wallis test, *P* = 1.1e-05) (Extended Data Fig. 11c). Another critical innate immune cell, neutrophils, were upregulated in SSC-III (Kruskal-Wallis test, *P* = 1.4e-05), Extended Data Fig. 11c). In terms of adaptive immunity, the results of the two methods (CIBERSORT and MCPcounter) diverged, but T cell abundance was slightly higher in SSC-III (Kruskal-Wallis test, *P* = 0.016) (Extended Data Fig. 11c). In general, the CD8+ cytotoxic T cell infiltration rate was extremely low for all three subtypes (Extended Data Fig. 11b-c). Taken together, SSC-II exhibited the highest stromal cell infiltration, whereas SSC-III was more infiltrated by innate immune cells, but less by adaptive immunity cells (Extended Data Fig. 11a-c). Additionally, we examined the expression levels of 59 immune checkpoints among SSCs, including 23 inhibitory and 36 stimulatory genes. As illustrated in Fig. 2b. SSC-III had critical inhibitory immune checkpoint overexpression (i.e., *PDCD1* (*PD-1*) and *CD274* (*PD-L1*)) (Kruskal-Wallis test, *P* = 6.5e-05 and *P* = 0.009, respectively). Of those, by applying intergroup analysis, *PD-L1* and *PD-1* expressions were significantly higher in SSC-III than SSC-I (Mann-Whitney test, *P* = 0.012 and *P* = 0.0023, respectively) and SSC-II (Mann-Whitney test, *P* = 0.0033 and *P* = 4.7e-05, respectively) (Fig. 2e). Subsequently, we implemented the TIDE algorithm to evaluate the potential clinical effects of immunotherapy in the three subtypes we defined. There was an apparent difference in the TIDE scores of the three SSCs. SSC-III had lower T cell dysfunction score than SSC-I (Mann-Whitney test, *P* = 2.6e-04) and SSC-II (Mann-Whitney test, *P* = 0.0011) and lower TIDE score than SSC-I (Mann-Whitney test, *P* = 1.4e-04) and SSC-II (Mann-Whitney test, *P* = 0.023), but the highest score regarding T cell exclusion (Mann-Whitney test, *P* = 4.2e-05 and *P* = 0.0021, respectively) (Extended Data Fig. 11d). SS homogeneously expresses multiple immunogenic cancer-testis antigens (CTAs) [34–37], predominately the CTA New York esophageal squamous cell carcinoma-1 (NY-ESO-1), the human melanoma antigen A4 (MAGEA4) as well as the preferentially expressed antigen in melanoma (PRAME). However, no significant differences in MAGEA4, NY-ESO-1 and PRAME could be observed across three SSCs (Extended Data Fig. 11e). Hence, these results may indicate an immune evasion phenotype of SSC-III and patients with this subtype may be more sensitive than other subtypes to anti-PD-1 or anti-PD-L1 therapies.

At the proteomic level, of the 13 samples that were feasible, 2 belonged to SSC-III, 7 to SSC-I and 4 to SSC-II. UMAP analysis of these 13 samples showed clear separation between SSC-III and SSC-I/SSC-II (Extended Data Fig. 11f). By comparing the expression level between DEGs and Differential expressed proteins (DEPs), we observed significantly strong correlations in the comparison of SSC-I to SSC-III (ρ = 0.556, *P* = 5.8e-191) and of SSC-II to SSC-III (ρ = 0.598, *P* = 3.4e-236) (Extended Data Fig. 11g). As expected, SSC-I was significantly enriched in cell cycle related pathways (Fig. 2f) and SSC-III (Fig. 2g) was significantly enriched in innate immune system related pathways, consistent with the results of the enrichment analysis in the RNA level.

Collectively, given the less distinguishable biological features between metastatic and primary tumors, our data suggested the gene expression patterns are determined *de novo* at the origin of the tumor. The subtype classification can well reflect biological features and reveal the heterogeneity of SS.

### Genomic characteristics of SS

Somatic mutations were detected in 39 out of 53 patients, with a median of one SNV per sample (Extended Data Fig. 12a). Most SNVs were missense mutations (Extended Data Fig. 12a). The most frequently mutated gene was KMT2C (21%, mutation frequency), albeit with low variant-allele frequency arguing against an important clonal driver function (Fig. 3a). Other recurrently mutated genes included *NOTCH1* (15%), *TP53*, *CTNB1*, *ATM,* or *HOXB13* (all at ≤5% frequency) (Extended Data Fig. 12a; Fig. 3a). Consistent with previous studies [38, 39], TMB in SS was characterized as very low, and in the pan-cancer setting, the TMB of SS was the lowest (per/50 MB) among the 33 cancer types (Extended Data Fig. 12b). Even though the TMB was identified to be low in SS, we observed different mutational events across SSCs. As shown in Fig. 3b, SSC-III had a higher TMB compared to SSC-II (Mann-Whitney test, *P* = 0.0033). Most cancer-related and putative functional genes were frequently mutated in SSC-III and I (Fig. 3c-d). In terms of functional analysis of these mutated genes, we explored activated pathways, and observed that NOTCH signaling was significantly enriched in SSC-III and I. However, gene alterations in the most enriched pathways (e.g., NOTCH signaling, Genome integrity, and Chromatin SWI/SNF complex) were not distinguishable between groups (metastasis and subtypes) (Fig. 3c). Interestingly, we found no difference in mutations between primary and metastatic lesions and no association between the presence of mutations and overall patient survival (Extended Data Fig. 12c-d). However, our cohort size might well be too small in order to detect any significant difference in outcome.

**Figure 3:**
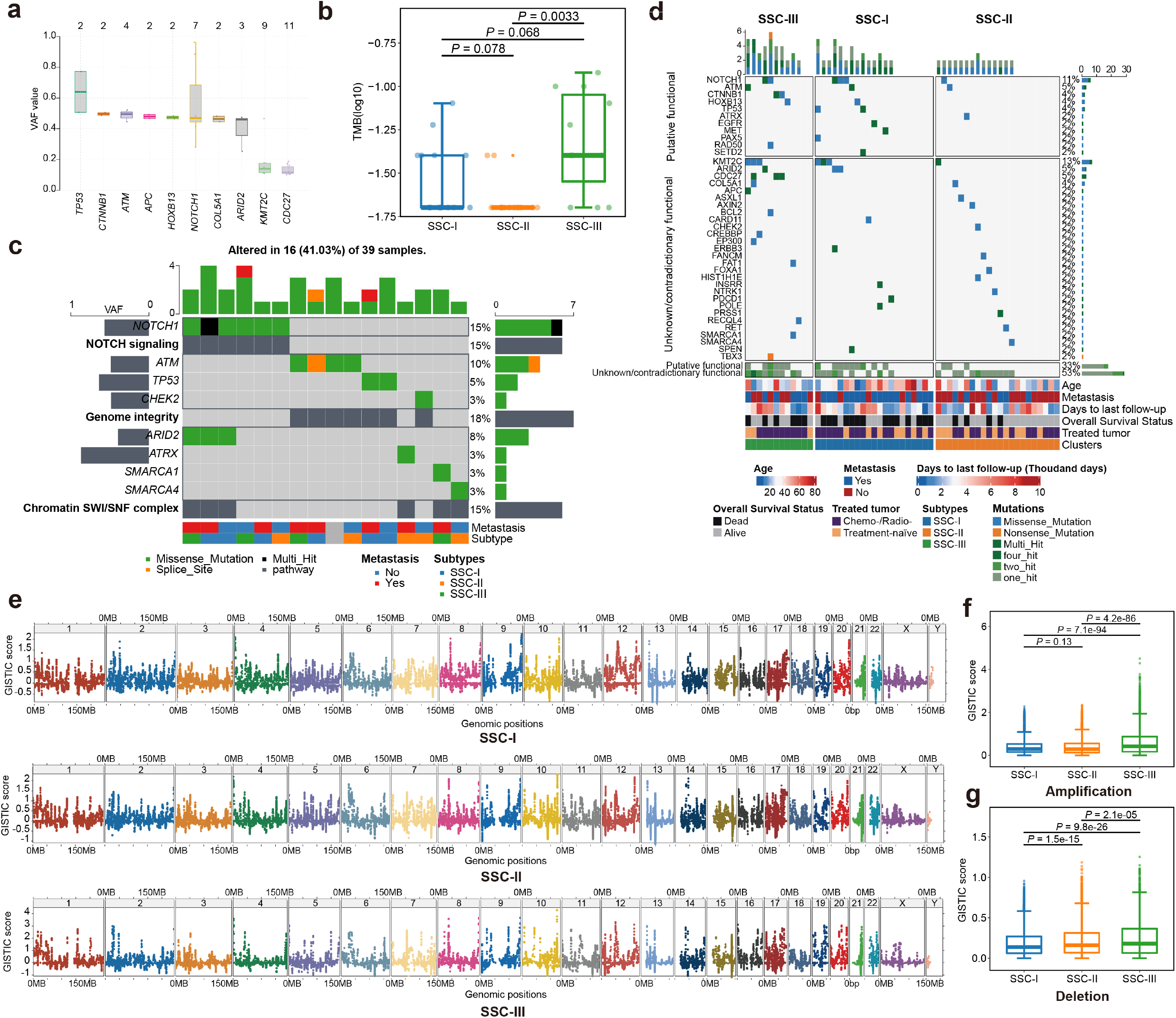
Genomic landscape of three SSCs. **(a).** Distributions of variant allele frequency in mutated genes. Middle line: median; box edges: 25th and 75th percentiles. **(b).** Distributions of the tumor mutation burdens (per 50MB, logarithmic scale) in the three SSCs. Middle line: median; box edges: 25th and 75th percentiles. Mann-Whitney U test. **(c).** Top three enrichment pathways of mutated genes and annotated below with metastasis and subtypes. **(d).** 55 SS patients with/without mutation data are ordered by their mutation frequencies and separated by subtypes, with clinical and molecular features annotations below. **(f).** Chromosome scatter plots depicting the GISTIC scores of three SSCs. **(g-h).** GISTIC scores distributions of **(g)** amplification and **(h)** deletions in the three SSCs. Middle line: median; box edges: 25th and 75th percentiles. Mann-Whitney U test.

To assess the somatic copy number alterations (SCNAs) that promote metastasis in SS, we applied an established approach (GISTIC 2.0) to calculate the SCNAs positive selection score at each genomic location. We found significantly higher frequency of amplification and deletions of genome-wide profiles of SCNAs in SSC-III as compared to SSC-I and SSC-II (Mann-Whitney test, *P* < 0.001) (Fig. 3e-g). Unexpectedly, we found a higher frequency of amplification in the genome-wide profiles of SCNAs in primary tumors than metastatic lesions (all sequenced samples) (Mann-Whitney test, *P* = 3.2e-04) (Extended Data Fig. 12e-f), and the GISTIC scores for both amplifications and deletions were significantly higher in patients without metastasis (all patients) (Mann-Whitney test, *P* = 2.9e-04 and *P* = 2.6e-20, respectively) (Extended Data Fig. 12g-h). These data suggest that there may be a clonal diversity in primary tumors and that metastatic lesions have undergone a clonal selection.

### Pre- and post-treatment effects of the expression and genetic features of SS

To comprehensively understand the correlation between cell morphology and immune infiltration, as well as the mechanisms of tumor evolution and therapeutic efficacy of SSCs, we obtained the droplet-based scRNA-seq data [22] to profile the malignant, immune, and stromal cells for the further investigation and validation of our bulk-sequencing results. We assigned cells to different cell types according to the DEGs and canonical markers (see Methods) (Extended Data Fig. 13a-b). We subdivided the malignant cells into three subsets, with canonical mesenchymal cycling (*TOP2A* and *MKI67*), mesenchymal (*SNAI2*, *PDGFRA*, *ZEB2*, and *ZEB1*), and epithelial (*MUC1* and *EPCAM*) markers[22, 40] to reflect the characteristics of SSC-I, SSC-II, and SSC-III, respectively. The other non-malignant subsets were B cells, CAFs, CD4 cells, CD8 cells, endothelial, mast cells, myeloid, NK cells, and pericytes (Extended Data Fig. 13c). Next, to be consistent with our cohort, only BP and MP patients were selected for downstream analysis (Fig. 4a). Interestingly, all patients but SS12pt had no T cell infiltrations (Fig. 4b).

**Figure 4:**
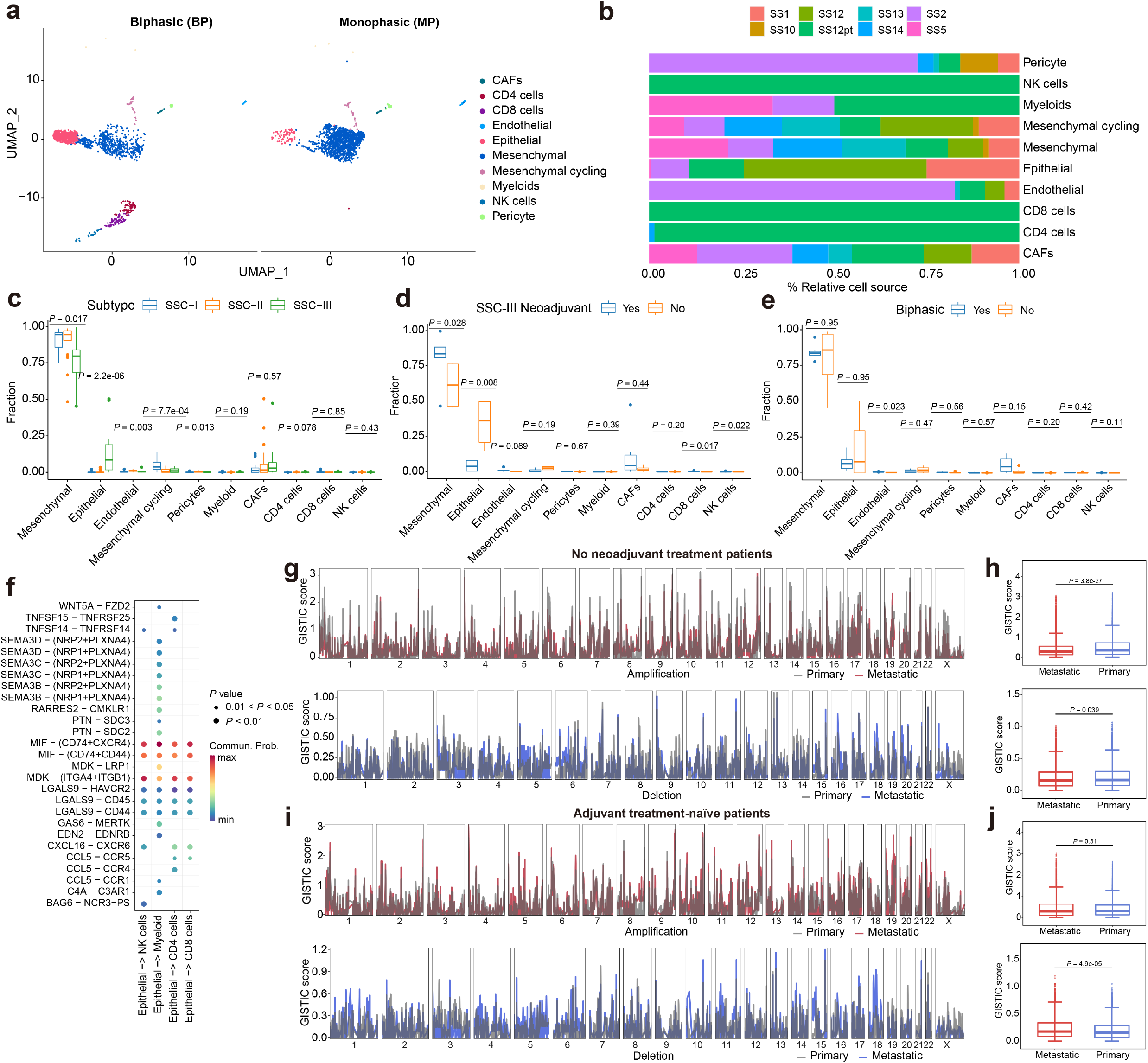
Treatment effects of SS. **(a).** UMAP plot of scRNA-seq profiles in BP and MP tumors, colored by cell types. **(b).** Distributions of the cell proportions in each sample. **(c).** The distributions of cell types in three SSCs after CIBERSORTx deconvolution. Middle line: median; box edges: 25th and 75th percentiles. Kruskal-Wallis test. **(d).** Cell type distributions in neoadjuvant and no neoadjuvant treated SSC-III patients after CIBERSORTx deconvolution. Middle line: median; box edges: 25th and 75th percentiles. Mann-Whitney U test. **(e).** Cell type distributions in neoadjuvant and no neoadjuvant treated biphasic patients after CIBERSORTx deconvolution. Middle line: median; box edges: 25th and 75th percentiles. Mann-Whitney U test. **(f).** Ligand–receptor pairs between epithelial cells and immune cells (NK cells, myeloid cells, CD4 cells, and CD8 cells. **(g).** GISTIC amplification (top) and deletion (bottom) plots of no neoadjuvant treatment patients. **(h).** GISTIC scores distributions of amplifications and deletions between primary and metastatic in no neoadjuvant treatment patients. Middle line: median; box edges: 25th and 75th percentiles. Mann-Whitney U test. **(i).** GISTIC amplification (top) and deletion (bottom) plots of adjuvant treatment-naïve patients. **(h).** GISTIC scores distributions of amplifications and deletions between primary and metastatic in adjuvant treatment-naïve patients. Middle line: median; box edges: 25th and 75th percentiles. Mann-Whitney U test.

Since the deconvolution approaches use different gene signatures that are widely utilized in many disease contexts, and since SS biology is distinct from other cancer types, we created a SS-specific gene signature to determine the TME of SS. By applying CIBERSORTx method [41], we generated the SS-specific gene signature based on the SS scRNA-seq data, and the new cell fraction of bulk RNA-seq data was re-imputed. As shown in Fig. 4c, SSC-II exhibited the highest mesenchymal and endothelial cell fractions (Kruskal-Wallis test, *P* = 0.017 and *P* = 0.003, respectively), SSC-III exhibited the highest epithelial cell fractions (Kruskal-Wallis test, *P* = 2.2e-06), and SSC-I exhibited the highest mesenchymal cycling cell fractions (Kruskal-Wallis test, *P* = 7.7e-04). The non-malignant cell infiltration was considerably lower than the presence of SS cells. Furthermore, we noticed that for patients with neoadjuvant treatment in SSC-III, the proportion of mesenchymal cells was significantly increased relative to pre-treatment patients (Mann-Whitney test, *P* = 0.028), whereas the epithelial cells were significantly decreased (Mann-Whitney test, *P* = 0.008) (Fig. 4c-e). However, malignant cell composition did not vary over treatment for the MP based subtypes (SSC-I and SSC-II) (Extended Data Fig. 13d-e) (Mann-Whitney test, *P* > 0.05) or for all BP patients in our cohort regardless of the treatment (Mann-Whitney test, *P* > 0.05) (Fig. 4e), suggesting that sensitivity of epithelial cells to treatment may be specific to SSC-III and controlled by other genetic events rather than inherent to the epithelium. Next, we examined the ligand-receptor interactions between malignant cells and infiltrating T-and myeloid cells. All these ligand–receptor pairs were expressed individually. Specifically, epithelial and mesenchymal cycling cells highly expressed the ligand Macrophage Migration Inhibitory Factor (MIF) and its receptors CD74, CXCR4, and CD44 in the immune cells (Fig. 4f; Extended Data Fig. 13f-g). In addition, MDK-(ITGAF+ITGB1) and CXCL12-CXCR4 showed a high probability of communication between epithelial and mesenchymal cells and immune cells, respectively. Since all these pathways are associated with immunosuppression, the results suggested an immune-resistant nature of SS, regardless of the malignant cell type.

In terms of the treatment effects in the genomic level, adjuvant therapy-naïve tumors had a slightly lower frequency of mutations compared to treated lesions (Extended Data Fig. 14), indicating that radiotherapy or chemotherapy may induce occasional mutations or trigger clonal selection in SS. When comparing tumor SCNAs in no neoadjuvant treatment patients, the frequency of copy number amplification and deletion was significantly higher in patients with primary tumors (Mann-Whitney test, *P* = 3.8e-27 and *P* = 0.039, respectively) (Fig. 4g-h). However, the frequency of deletion was significantly higher in all adjuvant therapy-naïve patients with metastases (Mann-Whitney test, *P* = 4.9e-05) (Fig. 4i-j), indicating that treatment resistant subclones may have a higher risk of developing metastasis which exhibits genomic stability.

### The regulation of mesenchymal to epithelial transition (MET) in SS

In order to further understand the biphasic SS biology, we sought to systematically explore the mechanisms that control the MET process. Through the trajectory plots, we observed a substantial number of transitions from mesenchymal to epithelial cells in BP patients, but these were considerably less noticeable in MP patients (Fig. 5a-b). Next, to identify the key genes involved in the MET process, a total of six overlapping genes were acquired among oncogenic program upregulated genes, Bulk RNA-seq DEGs (SSC-III vs SSC-II), scRNA-seq DEGs (BP-epithelial vs MP-epithelial), and scRNA-seq DEGs (epithelial vs other cells) (Fig. 5c). We observed that all these key genes, including *KIF1A*, *KRT8*, *CLDN4*, *KRT14*, *LY6E* and *FGF19* were overexpressed in BP epithelial cells, followed by the MET process (Extended Data Fig. S15a-b).

**Figure 5:**
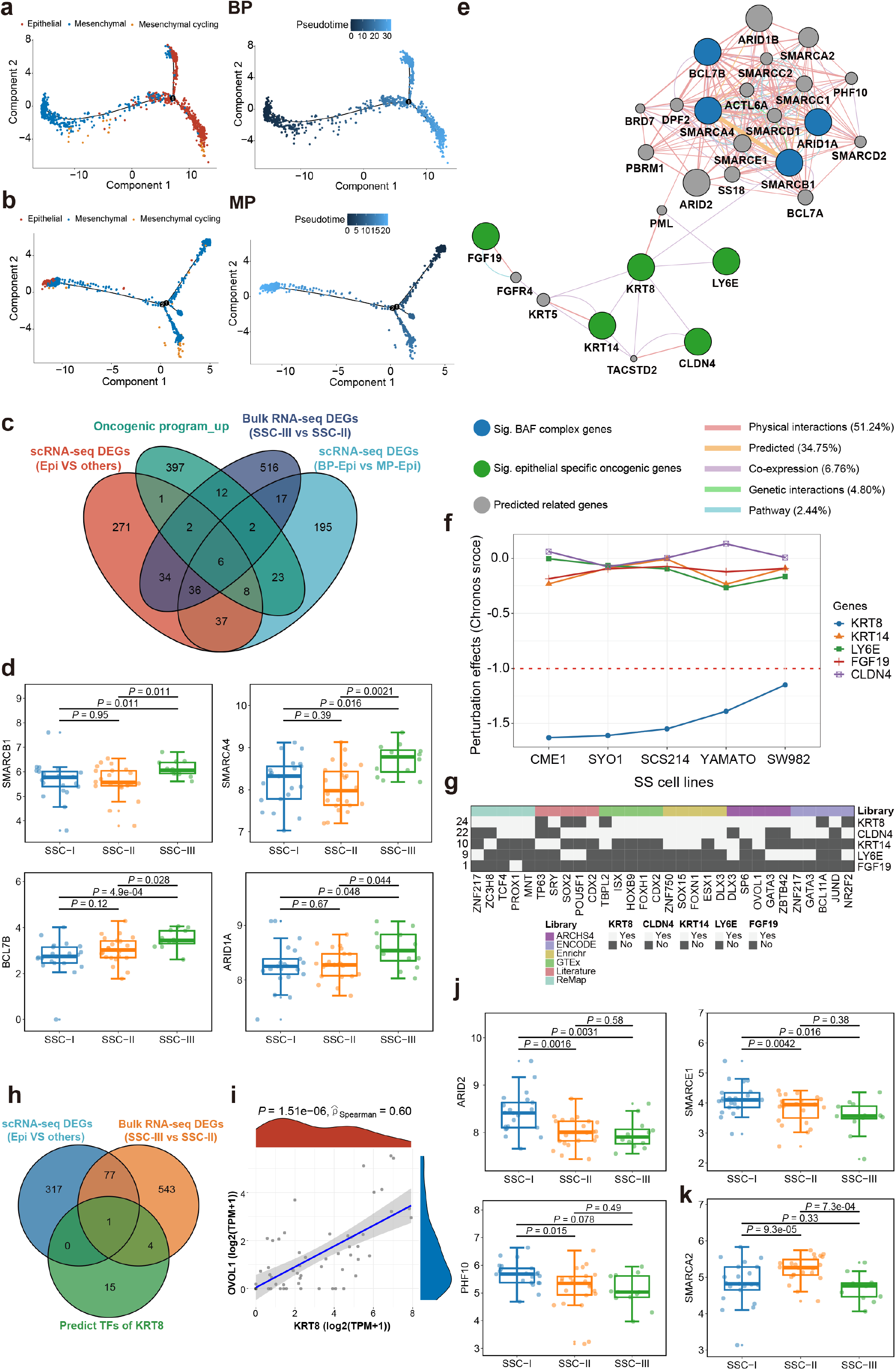
The regulation of mesenchymal to epithelial transitions in SS. **(a-b).** The trajectory plots showing the dynamic development of epithelial, mesenchymal, and mesenchymal cycling cells in **(a)** BP and **(b)** MP patients and their pseudotime curve. **(c).** Venn diagram depicting the overlap of the MET key genes involved in oncogenic program upregulated genes, Bulk RNA-seq DEGs (SSC-III vs SSC-II), scRNA-seq DEGs (BP-epithelial vs MP-epithelial), and scRNA-seq DEGs (epithelial vs other cells). **(d).** The gene expression distributions of *SMARCB1*, *SMARCA4*, *BCL7B*, and *ARID1A* across SSCs. Middle line: median; box edges: 25th and 75th percentiles. Mann-Whitney U test. **(e).** The predicted gene interaction network between significant BAF complex genes, epithelial specific oncogenic genes, and other putative additional relevant genes. **(f).** Perturbation effects of knockdown MET key genes in five SS cell lines. **(g**). Schematic overview of predicted TFs of MET key genes **(h).** Venn diagram depicting the overlap of the MET key genes involved in predicted TFs of *KRT8*, Bulk RNA-seq DEGs (SSC-III vs SSC-II), and scRNA-seq DEGs (epithelial vs other cells) **(i).** The correlation between expression levels of *KRT8* and *OVOL1*, Spearman rank correlation. **(j).** The gene expression distributions of *ARID2*, *SMARCE1*, *PHF10*, and *SMARCA2* across SSCs. Middle line: median; box edges: 25th and 75th percentiles. Mann-Whitney U test.

Since the chimeric oncoprotein SS18-SSX is considered as the primary driver and hallmark genetic alternation of SS and involved in chromatin remodeling for mediating sarcomagenesis [9, 10]. Specifically, the chromatin remodeling switching defective (SWI)/ sucrose non-fermenting (SNF) family of multiprotein complexes are critical to recruiting additional proteins for chromatin remodeling and transcriptional regulation in SS, including canonical BAF (cBAF), polybromo-associated BAF (pBAF), and noncanonical BAF (ncBAF) [42–45]. To examine the gene expression changes of the BAF complex subunits across three SSCs, we found that BAF complex subunits, including *SMARCB1*, *SMARCA4*, *BCL7B*, and cBAF complex specific subunit *ARID1A* were significantly upregulated in SSC-III (Fig. 5d). A network of the significant BAF complex genes and MET key genes was constructed (Fig. 5e). Interestingly, *KRT8* was implicated in MET and has a potential direct interaction with the BAF complex, whereas other key genes (e.g., *LY6E*, *CLDN4*, *KRT14*, and *FGF19*) were acting as the downstream regulators of this network. Furthermore, according to the data on the perturbation effect of the Dependency Map (DepMap) portal [46, 47], only *KRT8* had the lowest Chronos score among the five SS cell lines, indicating an essential gene in SS cell lines required for survival (Fig. 5f). To investigate the potential regulatory mechanisms of *KRT8* expression in SS, 24 distinct transcription factors (four overlaps) regulating *KRT8* were predicted based on six libraries using the ChEA3 database [48] (Fig. 5g). As a result, only one TF, *OVOL1*, was overlapped among predicted TFs, bulk RNA-seq DEGs (SSC-III vs SSC-II), and scRNA-seq DEGs (epithelial vs other cells) (Fig. 5h). As expected, there was a high correlation of gene expression levels between *OVOL1* and *KRT8* (ρ = 0.60, *P* = 1.51e-06). Taken together, *KRT8* may play a critical role in the MET process of biphasic SS biology and could be regulated by TF *OVOL1*. Moreover, the pBAF complex subunits ARID2, SMARCE1, and PHF10 were significantly overexpressed in SSC-I (Mann-Whitney test, *P* < 0.05) (Fig. 5j), whereas the common BAF complex ATPase SMARCA2 was significantly overexpressed in SSC-II (Mann-Whitney test, *P* < 0.05) (Fig. 5k).

### Identification of SCNA drivers in SS

To further investigate the potential drivers facilitating the treatment and to understand the biology of SSC-III in terms of the low proliferation and high immune response, we computed the genome-wide *q*-value for identifying the differential chromosome arm-level copy-number events by comparing to SSC-II (GISTIC score difference > 1, *q*-value < 0.001) (Fig. 6a-c). By identifying the key drivers in both genomic and transcriptomic levels, we overlapped the significantly amplified genes, the gene panel for the target sequencing, and the DEGs between SSC-III and SSC-II. As a result, two genes), the *PD1* and *TMPRSS2*, located in the 2q37.3 regions were significantly up-regulated in SSC-III (Fig. 6e (q-value = 4.981e-08) and 21q22.3 (q-value = 4.7977e-07) respectively. To understand these biological contexts of these amplifications, we examined the prognostic impact of *PD1* and *TMPRSS2* high amplification status. As shown in Fig. 6f-g, in SSC-III, patients with *PD1* or *TPMPRSS2* had better outcome than the ones without. PD1 + TMPRSS2 high amplification status had the highest AUC of metastasis/death (AUC = 0.833), followed by *TMPRSS2* (AUC = 0.800), *PD1* (AUC = 0.733), and neoadjuvant response (AUC = 0.667).

**Figure 6:**
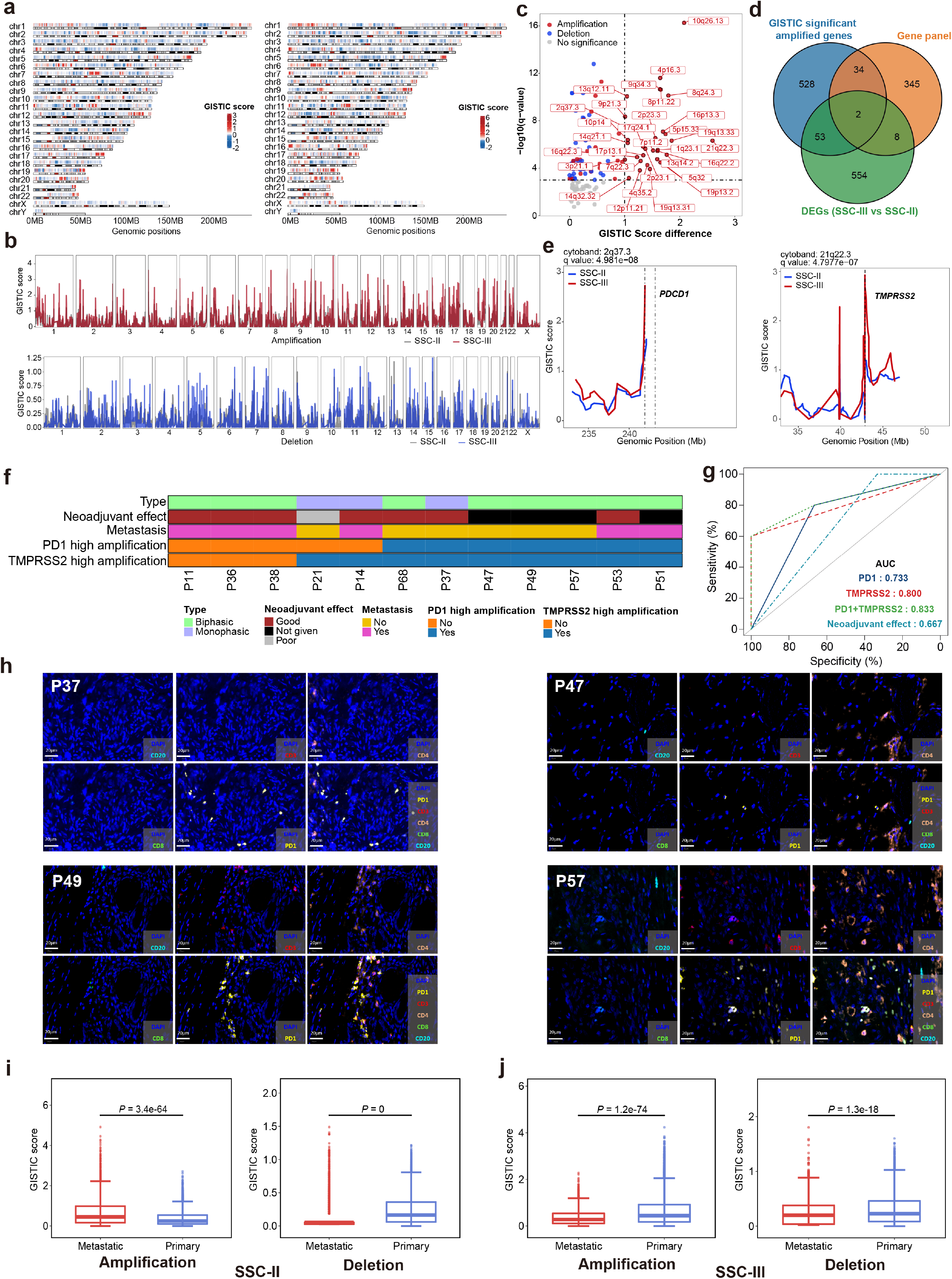
The secondary driver SCNAs in biphasic SS. **(a).** Chromosome heatmaps depicting the GISTIC scores in chromosome 1-22, X, and Y. **(b).** GISTIC amplification (top) and deletion (bottom) plots of SSC-II and SSC-III. **(c).** Differentially amplified regions in SSC-III compared to SSC-II. Significant differential regions are labeled (*q*-value < 0.001; GISTIC score difference > 1). **(d).** Venn diagram depicting the overlap of the candidate drivers involved in significant amplified genes, DEGs (SSC-III vs SSC-II), and gene panel (389) genes. **(e).** GISTIC plots of differential region and candidate driver regions. **(f).** Clinical characteristics and the high amplification status of *PDCD1* and TMPRSS2 in SSC-III patients. **(g).** ROC curves of the predication performances *PDCD1*, TMPRSS2, and neoadjuvant effect in MFS. **(h).** Opal multiplexing visualizing various markers in synovial sarcoma tissue. Blue pseudocolor = nuclei; aqua = CD20; red = CD3; orange = CD4; green = CD8; yellow = PD1. **(i-j).** GISTIC score distributions of amplifications and deletions between primary and metastatic in **(i)** SSC-II and **(j)** SSC-III samples. Middle line: median; box edges: 25th and 75th percentiles. Mann-Whitney U test.

Four patients (P37, P47, P49, and P57) with both *PD1* and *TMPRSS2* high amplification status were subjected to multiplex IHC staining (Extended Data Fig. 16a-b). As shown in Fig 6h, these patients all had low immune cell infiltration and a certain amount of PD-1 expression in the tumor cells, which were consistent with the findings based on the RNA and DNA data. To project these phenomena into all biphasic patients, we found that high PD1 amplification was significantly associated with KRAS signaling downregulation (Mann-Whitney test, *P* = 0.0027) (Extended Data Fig. 16c). Regarding the functional enrichment of *TMPRSS2* high amplification, it was significantly associated with certain signaling pathways, including estrogen early response, MTORC1 signaling downregulation, and KRAS signaling downregulation (Mann-Whitney test, *P* = 0.029, *P* = 0.013, and *P* = 0.043, respectively) (Extended Data Fig. 16d); lower proliferation, including E2F targets, MYC targets V1, G2M checkpoint, and P53 pathway (Mann-Whitney test, *P* = 0.02, *P* = 0.008, *P* = 0.02, and *P* = 0.043, respectively) (Extended Data Fig. 16e); lower metabolism, including adipogenesis, oxidative phosphorylation, bile acid metabolism, and fatty acid metabolism (Mann-Whitney test, *P* = 0.013, *P* = 0.029, *P* = 0.008, and *P* = 0.02, respectively) (Extended Data Fig. 16f). In addition, BP patients with *TMPRSS2* high amplification had slightly higher immune responses (Extended Data Fig. 16g). Taken together, *PD1* and *TMPRSS2* high amplification status could be regarded as favorable predictors for biphasic patients. The favorable prognostic impact of tumor-intrinsic PD1 following genetic amplification might contribute to ineffectiveness of anti-PD1 immunotherapy in this patient group containing the most highly immune-cell infiltrated SS patients.

Next, we performed longitudinal analyses for all treated tumors. Metastatic tumors displayed significantly higher amplification levels than primary tumors in SSC-II (Mann-Whitney test, *P* = 3.4e-64) (Fig. 6i), but lower in SSC-III (Mann-Whitney test, *P* = 1.2e-74) (Fig. 6j). However, in SSC-I, the difference was less significant (Mann-Whitney test, *P* = 0.03) (Extended Data Fig. 16h). Among all MP tumors, metastatic tumors had higher deletion levels (Mann-Whitney test, *P* = 2.8e-18), while among all BP tumors, primary tumors had higher SCNA regarding both amplifications and deletions (Mann-Whitney test, *P* = 9.8e-19 and *P* = 2.8e-229, respectively) (Extended Data Fig. 16i-j). To conclude, treatment-sensitive BP tumors harbor more SCNAs, while *PD1* and *TMPRSS2* could be regarded as secondary drivers of BP SS.

Herein, we proposed a tumor evolution model for SS. As illustrated in Fig.7, pathological chromosomal translocations t(X;18)(p11;q11) occur in normal cells and induce epigenetic dysregulations by remodeling the BAF complex. Preexisting or acquired differences in the BAF complex composition led to different epigenetic effects, resulting in three major branches of tumor evolution in SS, including i). Hyperproliferative-SS (monophasic), ii). Vascular-SS (monophasic), and iii). Epithelial-SS. Immune resistance appears to occur *de novo*. Hyperproliferative-SS may primarily be regulated by pBAF complex-based epigenetic modifications, resulting in overexpression of highly oncogenic program-cycling genes and high therapeutic resistance and E2F target genes, which can be treated by drug combinations of HDAC and CDK4/6 inhibitors [22]. Vascularized-SS may primarily be regulated by SMARCA2-based epigenetic modifications, and its metastasis may be accelerated by secondary mutations or SCNAs. Epithelial-SS may primarily be regulated by cBAF complex-based epigenetic modifications, leading to an early MET. While the high SCNVs could cause the malignant epithelial cells to harbor more treatment sensitive clones, the mesenchymal cells still have substantial chemotherapy resistance with high rates of recurrence and progression.

**Figure 7:**
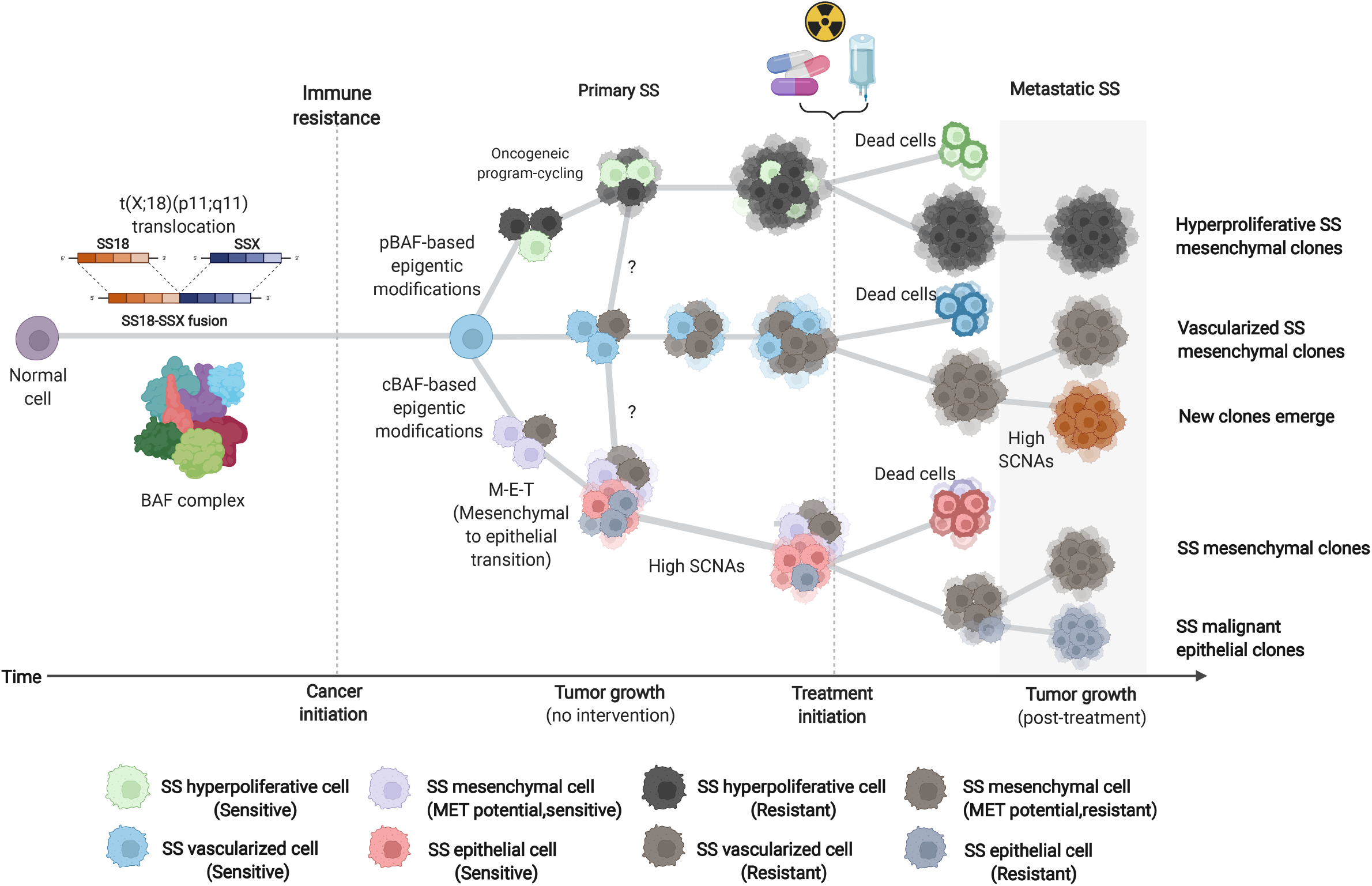
The tumor evolution model of SS. Schematic illustration of SS tumor evolution in cancer initiation, tumor development, and treatment interventions. Round and irregular cells represent normal cells and cancer cells, respectively. The light and dark colors represent different SS-sensitive and resistant cells, respectively. The size of tumor masses represents estimated different cell proportions. Cell membrane thickness represents necrotic cells (thick) and normal growing cells (thin).

## Discussion

Herein we present an integrated analysis of transcriptome sequencing, targeted DNA sequencing, and LC-MS proteomics of SS. In line with previous studies, we found low levels of secondary mutations, and varying levels of chromosome complexity. Not surprisingly, therapy naïve tumors that developed metastasis had higher levels of SCNAs compared to tumors that did not develop metastasis, indicating that SCNAs might be important for tumor progression. Interestingly we observed the opposite relationship in tumors that had been treated with chemotherapy, indicating that primary tumors may contain multiple clones and post-treatment metastasis have less complex SCNAs indicating a clonal selection during therapy. We found that a substantial proportion of cases harbored mutually exclusive mutations in genes associated with genome integrity (18%) or the BAF complex (15%), but we were unable to find any association to clinical outcome or molecular phenotype. We also identified and validated non-recurrent structural variants resulting in secondary fusion genes. While most of the validated fusion genes were predicted to produce functional proteins, we were unable to provide evidence for their role as oncogenic drivers. Since we were unable to identify any clinically relevant secondary fusion genes, we conclude that they likely constitute stochastic events with limited biological relevance in SS.

We identified three distinct molecular patterns which were largely similar to the histological subtypes of SS. Based on the transcriptome patterns we named these clusters SSC-I: Hyperproliferative-SS (resembles poorly-differentiated SS); SSC-II: Vascularized-SS (similar to monophasic SS); and SSC-III: Epithelial-SS (similar to biphasic SS).

Patients with hyperproliferative-SS had the worst prognosis. These tumors were distinguished by high expression of E2F and MYC targets functionally related to proliferation, DNA repair pathways and the previously described *core oncogenic program* [49]. SS18-SSX may induce degradation of the canonical BAF-complex (cBAF). Since we observed significantly higher expression of the polybromo-associated BAF (pBAF) specific components *ARID2*, *SMARCE1* and *PHF10* in SSC-I, it is implied that SSC-I is dominated by the pBAF complex [50, 51]. SCNAs were similar to vascularized SS (SSC-II) and evidently less frequent than in epithelial-SS (SSC-III), indicating that the hyperproliferative phenotype arises *de novo* and is not a progression from SSC-II/III. This was also supported by the absence of switching between SSCs in patients with multiple sequenced lesions at different time points. The observations of clonally evolutionary distinct subgroups are in line with the observations done by Przybyl *et al.* [52]. Interestingly, many tumors in the hyperproliferative-SS group were not histologically characterized as poorly differentiated SS, suggesting that the histological criteria for classifying poorly differentiated SS should be reviewed with this in mind.

The vascularized-SS (SSC-II) cluster was characterized by better patient outcome (the 10- and 20-year overall survival was 75%). These tumors had more non-tumor stroma, higher expression of vascular-related genes, downregulation of DNA-repair pathways and the lowest expression of the *core oncogenic program*. While most vascularized-SS had a very low tumor mutational burden and SCNA alterations, patients that died of disease in this group had distinctly higher frequency of SCNA amplifications (Fig. 1g) indicating that second hits might be required for metastasis in SCC-I. Although this group had better outcome it would be hazardous to give any recommendations regarding therapy and grading, since it is not clear whether the outcome is related to better sensitivity to chemotherapy, a less malignant phenotype or a combination of the two.

The epithelial-SS (SSC-III) was dominated by biphasic tumors and characterized by higher levels of kinase-signaling targets, innate immune cell infiltration and lower levels of proliferation and DNA repair. These tumors had significantly higher expression of the BAF subunits *SMARCB1*, *SMARCA4*, *BCL7B*, and cBAF complex specific subunit *ARID1A* suggesting a relatively higher retention of cBAF. Since this group had the highest levels of SCNA and somatic mutations it is natural to assume that these genomic alterations constitute defining drivers for SSC-III. We performed expression network analysis and identified a low molecular weight cytokeratin (KRT8) as a pivotal gene for MET in SS. KRT8 can interact with the BAF complex, and plays a role in maintaining cellular structural integrity and functions in signal transduction and cellular differentiation[53]. We also predicted that the key to KRT8 regulation would be the OVOL1 transcriptional factor, which has been shown critically regulate the differentiation of epithelial cells [54, 55]. However, further work would be required in order to understand the regulatory mechanisms transform SS cells to an epithelial phenotype. Surprisingly, while many SSC-III showed a good histological response to neoadjuvant chemotherapy (<10% viable tumor cells), the long-term MFS and OS were both very poor. We were able to identify that patients whose tumors had a higher frequency of CNA amplifications (including *PD1*, *TMPRSS2*) had a more favorable prognosis after neoadjuvant therapy, indicating that chemosensitivity could be identified through a stratification algorithm.

The differences in immunological signatures between SSCs prompted us to further investigate immune-related patterns and immune cell compositions. Petitprez et al. proposed five sarcoma immune classes, where half of the synovial sarcoma cases were classified in an immune-low *immune desert* phenotype, while the remaining belonged to *immune-high* or *highly vascularized* groups [56] which show striking similarity with our SSC classification. Compared to other forms of cancer, SS is characterized by a low adaptive immune infiltration and exhibits a considerably high infiltration of naïve and quiescent B cells and dysfunctional CD8^+^ T cells. Following cell composition deconvolution using SS-specific gene signatures, we found limited differences in immune cell infiltrations between SSCs, but all groups demonstrated evidence of immune suppression.

The interplay between the oncogenic process and tumor immune evasion has been previously described in SS. A combination of HDAC and CDK4/6 inhibitors targeted the malignant state of SS cells and increased their immunogenicity, thereby augmenting T cell reactivity and T cell-mediated killing capacity [22]. Our transcriptome analysis identified that both hyperproliferative-SS (SSC-I) and epithelial-SS (SSC-III) with low CNVs had high enrichment of proliferative pathways and might benefit from this drug regimen. Most importantly, our data support the notion that molecular subtyping might be important for patient stratification in new treatment regimes.

In SSC-III we observed upregulated expression of PD-1 (and gene amplification) and PD-L1 which could indicate immune check-point activation. Recent studies have described non-canonical PD-1 signaling different tumor types. In malignant melanoma and pancreatic cancer, PD-1 signaling promotes cancer cell proliferation by interacting with mTOR and Hippo signaling [57, 58]. In non-small cell lung cancer (NSCLC) and colon cancer, PD-1 signaling inhibits cancer cell proliferation through inhibition of AKT and ERK signaling [59, 60]. By using multiplex IHC we confirmed that tumor cell expressed PD-1, and this was significantly associated with downregulation of KRAS signaling, indicating that the nature of PD-1 signaling in epithelial-SS is more similar to that observed in NSCLC and colon cancer. So far, it has been reported that CTLA-4 and PD-1 inhibitors failed to demonstrate clinically relevant benefits in synovial sarcoma with a response rate below 10% [25, 26, 61]. Given the rapid development of new therapeutic strategies with immune checkpoint inhibitors, these observations should provide a rationale for further investigating immunotherapy in a subset of SS.

Although the bioinformatics approaches for analyzing bulk RNA and DNA sequencing data have been well established, this study have some inherent limitations. First, our study lacked the number of cases to make robust longitudinal analyses to observe the difference in metastatic evolutions in the various SSCs. Second, the true predictive effect of the gene signature-based classification needs to be estimated in a prospective study. Finally, while we identified a potential role for immune checkpoint inhibitors in a subset of SS, we had no patients that had been treated with immunotherapy.

To conclude, our results revealed a substantial heterogeneity among SS and facilitated patient stratification into three subtypes based on multi-omics data analysis. We described the biological and genetic variability between the subtypes, including key transcriptome patterns, putative mechanistic differences related to the composition of the BAF-complex and secondary genomic events, including SCNA which could be associated with the long-term outcome after chemotherapy. All SS exhibited an immune inhibitory phenotype, albeit with some differences including expression of immune checkpoint markers. We believe that these results explain parts of the biology underlying SS and will help us understand the differences in clinical presentation which may prove important for future therapy development and patient stratification.

## Acknowledgments

The authors would like to acknowledge the expertise and support with NGS services provided by the National Genomics Infrastructure (NGI) and the Clinical Genomics Stockholm facility at the Science for Life Laboratory (jointly hosted by the Department of Microbiology, Tumor and Cell Biology at Karolinska Institutet and Department of Gene Technology at School of Engineering Sciences in Chemistry, Biotechnology and Health at KTH Royal Institute of Technology) and the Genomic Medicine Center Karolinska at the Karolinska University Hospital. The computations were enabled by resources in the project [sens2019015] and [SNIC2020/15-87] provided by the Swedish National Infrastructure for Computing (SNIC) at UPPMAX, partially funded by the Swedish Research Council through grant agreement no. 2018-05973.

## Author contributions

Y.C. and F.H., conceived the project, designed the study. Y.C., I.R.L., I.S., H.J., W.- K.H, J.L. and F.H. interpreted results. F.H. and O.L. obtained funding for the study. P.T., A.C.H., A.P., Y.Z. and F.H. consented patients for the study, collected synovial sarcoma samples, and provide the clinical data. Y.Z. and F.H. performed the pathological evaluations and tissue biopsies. Y.C. performed transcriptomic and genomic analyses. I.S. and Y.C. performed the proteomics analyses. J.Z. provided support for single-cell RNA analyses. Y.C. and Y.L. performed RNA and DNA sample preparations. Y.C., X.C., and H.J. performed proteomics sample preparation, MS data generation and searching. X.L. and Y.L. performed the experimental validations of the fusion genes. Y.S. and M.E. performed the multiplexing and digital image analysis. Y.C. and F.H. wrote the manuscript with feedback from all authors.

## Data availability

The processed RNA-seq data will be provided through the Gene Expression Omnibus (GEO); The processed panel sequencing data will be provided via the European Genome-Phenome Archive (EGA); The processed MS proteomics data will be deposited on the ProteomeXchange Consortium via the (Proteomics Identification Database) PRIDE partner repository.

## Materials and Methods

### Patient samples and ethics

The digital patient records of the Pathology Clinic at the Karolinska University Hospital were searched for cases of synovial sarcoma. Formalin-fixed paraffin-embedded (FFPE) tissue was collected from all cases classified as SS in the archive and was reviewed. In retrospect and without the detailed mapping of treatment protocol or pathology sampling protocol, primary tumors that had been treated with neoadjuvant treatment were classified as good responders when ≤10% of the tumor was deemed viable or as poor responders when >10% of the tumor was deemed viable. To include a case in the study, we required DNA or RNA with sufficient quality for sequencing and proof of the SS18-SSX fusion gene. Clinical FISH or RT-PCR was available for most cases, but to further validate the diagnosis, all cases were confirmed positive for the SS18-SSX chimeric-protein specific antibody. In the end, a total of 55 patients with SS were included in the study. Fifty patients had localized tumors, and five patients had metastatic disease at the time of diagnosis. All patients were surgically treated at the Karolinska University Hospital between 1992-2016, and the follow-up data collection was concluded in December 2019. The clinical information is shown in Supplementary Table 1.

The study and collection of patients’ samples were approved by the local ethics committee (The Regional Ethics Committee in Stockholm), reference number 2013 1979-31. All patients had given oral and written consent prior to sample collection. The study was performed in agreement with the Declaration of Helsinki.

### Publicly available gene expression data

For the validation cohorts, the microarray-based gene expression data of GSE40021(58 SS cases) [18] and single-cell RNA-seq (scRNA-seq)-based gene expression data of GSE131309 (12 SS cases) [22] were obtained from the NCBI Gene Expression Omnibus (GEO) database with corresponding clinical data. For the pan-cancer analysis, we downloaded the uniformly normalized pan-cancer dataset: TCGA Pan-Cancer (PANCAN, N=10535, G=60499) from the UCSC (https://xenabrowser.net/) database, the gene expression profile of each tumor was extracted separately and mapped to the genome annotation file (hg38). In addition, we performed log2 (x+1) transformation of each expression value. In terms of the genomic data, we obtained the Simple Nucleotide Variation dataset of the level4 of all TCGA samples processed by MuTect2 [62] software from GDC (https://portal.gdc.cancer.gov/), and calculated the tumor mutation burden (TMB) of each tumor using the “tmb” function of the R package “maftools” (v 2.6.05) [63], We also excluded cancer types with less than three samples, and finally obtained the expression data of 33 cancer types.

### Tissue DNA and RNA extraction

After histological review, areas with high tumor purity (range 95 −75%) were dissected from FFPE blocks using 1 mm punch biopsies. Tumor total RNA and DNA were extracted from 91 lesions, where 55 were from primary tumors and 36 from metastatic lesions. Normal DNA was extracted from adjacent normal tissue (n = 88). A summary of the sample processing is shown in Fig. 1a.

### Whole transcriptomic analysis

Library generation, quality control, sequencing, and initial data processing were performed at the National Genomics Infrastructure, Science for Life Laboratories in Stockholm. RNA library preparation was done using the Illumina TruSeq Stranded Total RNA and the Illumina RiboZero. Clustering was done by ‘cBot,’ and samples were sequenced on NovaSeq6000 (NovaSeq Control Software 1.6.0/RTA v3.4.4) with a 2×151 setup using ‘NovaSeqStandard’ workflow in ‘SP’ mode flowcell. The Bcl to FastQ conversion was performed using bcl2fastq_v2.19.1.403 from the CASAVA software suite. The quality scale used was Sanger / phred33 / Illumina 1.8. The average sequence depth was 37.5 M reads (min 16.4 – max 82.3 M reads).

### RNA seq data processing

RNAseq data were processed with the nf-core RNAseq pipeline v1.0[64]. Default parameters were used unless mentioned otherwise. Sequencing data quality was assessed using the FastQC [65]. TrimGalore was used to remove adapter contamination and trim low-quality areas (https://www.bioinformatics.babraham.ac.uk/projects/trim_galore/), with the Cutadapt function for adapter trimming and the FastQC function upon completion[66]. Sequences were aligned to the human reference genome (hg19/GRCh37) by applying the STAR software. Transcript counts were produced with featureCounts [67] using the transcriptome annotation file generated by StringTie [68]. We reported the gene expression data in both counts and transcripts per million (TPM) values. Raw counts were used for differential expression analyses (“DESeq2” package[69] in R). For quantification purposes, Transcripts per million (TPM) were calculated using Kallisto (version 0.46.1) [70]. Transcripts with a TPM score above one were retained, resulting in a total of 18934 gene IDs. All known exons in the annotated file were 100% covered. Log transformation of TPM values was calculated as log2 (TPM + 1) and applied for subsequent analyses.

### Fusion gene detection

Screening for chimeric fusion transcripts was performed on raw fastq files by two different bioinformatics pipelines, FusionCatcher [71] (Version 1.10) and STAR-Fusion [72] (Version 1.2.0) with standard settings. We conceived a series of filtering processes to retain the putative true-positive gene fusions as FusionCatcher and STAR-Fusion both reported multiple fusion genes per sample, many of which had a high probability of being false-positive [72, 73]. Since output files used different formats, we applied two different filtering processes. For FusionCatcher, we excluded fusion genes a). on a list of known false positives (banned); b). fusion genes mapped to multiple genomic locations indicative of sequence homology (high common mapping reads); c). if one fusion gene partner had at least nine other fusion gene partners (promiscuous genes); d). fusion genes with less than three unique sequencing reads (low bioinformatic support); e). if both genes forming the fusion were adjacent or closely located in the genome (adjacent); f). the sequence of the fusion junction contained a highly repetitive region containing repeating short sequences or polyA/C/G/T (short repeats). For STAR-fusion, we removed a). fusion genes with adjacent partners; b). fusions resulting from very local gene rearrangements; c). fusion transcripts with non-canonical splice site sequences; d). if one gene was fused with nine or more genes and e). genes that encoded procadherins. Subsequently, all filtered fusion genes were manually inspected using the Integrative genomics viewer (IGV) [74] and the R package “chimeraviz” [75] to visualize the aligned reads. All the fusion gene information is shown in Supplementary Table 2.

### Fusion gene verification using sanger sequencing

To validate the fusion genes detected by RNA-seq, RNA was isolated from the same tumor with a separate punch biopsy using the QIAamp RNeasy FFPE Extraction Kit (Qiagen, Cat # 73504). Sequencing primers were designed according to the full sequences and breakpoints using the output files from STAR-fusion and FusionCatcher. Each fusion gene was verified using two sets of primer pairs. cDNA was synthesized using High-Capacity RNA-to-cDNA^TM^ Kit from Thermo Scientific (Cat # 4387406). After touch-down PCR reactions (40 cycles), the amplification products were electrophoresed in a 5% agarose gel. The fusion genes were considered to pass the PCR validation if the size of the product corresponded to the expected size. The PCR products were isolated and purified by QIAquick PCR purification Kit (Qiagen, Cat # 28104), and TA cloned into pCR™4-TOPO® vector using TOPOTM Cloning TM Kit (Thermo Fisher, Cat # 450071). The vectors were transformed into DH5a *E. coli* and cultured overnight on an LB agar plate, whereupon clones were selected and identified by PCR for DNA insertion. For positive clones, amplification in LB medium with 100ug/ml AMPr was conducted overnight. Finally, vectors were purified with the QIAprep Spin Miniprep KitBody Text (Qiagen, Cat #27014) and Sanger sequenced with M13-primers (Eurofins Genomics, Cologne, Germany) and manually analyzed using the 4Peaks Software (version 1.7.1, Mekentosj). All the whitelist fusion gene sequences and primers are shown in Supplementary Table 3 (some whitelist fusion genes were excluded from PCR validation due to exhaustion of FFPE samples).

### Gene network construction and functional enrichment analysis

Both Gene Set Enrichment Analysis (GSEA v4.0) [76, 77] and Metascape [78] were used for pathway enrichment analysis. Specifically, samples of all gene expression data (normalized) were pooled according to the paired grouping of patients and were subjected to GSEA. Molecular Signatures Database [79] (MSigDB v7.4) of Hallmark gene sets (H), Reactome gene sets (C2), and Gene Ontology (GO) biological process gene sets (C5) were employed for enrichment analysis. Normalized *P*-value <0.01 and FDR < 0.01 were selected as the threshold as significance. Additionally, Metascape was used to identify gene enrichment terms in up-or down-regulated genes for GO and Reactome pathway analyses. To identify gene expression patterns associated with the metastatic disease, we performed five comparisons for GSEA: i) Comparison 1: all metastatic samples compared to all primary samples; ii) Comparison 2: Over follow-up time in 10 patients with multiple lesions, metastatic samples compared to all primary samples; iii) Comparison 3: patients with metastases versus without; iv) Comparison 4: treatment-naïve patients with metastases versus without; v) Comparison 5: patients with metastatic events versus without in a validation cohort-GSE40021). Analysis for gene networks and pathways was performed with the Cytoscape v3.8.2 software [80] with the GeneMANIA [81] plugin according to the instruction manual. We analyzed gene networks to identify gene-to-gene interactions, the topology of such gene correlations, and putative additional genes that may be involved in significant BAF complex genes and epithelial cell-specific oncogenes if shown to interact with the large number of genes in the query set.

### Immune profiling analyses

We performed seven different computational algorithms to assess the immune cell content in the tumor microenvironment (TME), and the results were confirmed by assessing the consistency of the abundance/fraction of immune cells in the cross-validation of deconvolution methods. We performed Cell-type identification by estimating relative subsets of RNA transcripts (CIBERSORT) [82] to quantify the fractions of the 22 immune cell subtypes. In terms of immune cell abundance, we utilized the Estimation of STromal and Immune cells in MAlignant Tumor tissues using the Expression data (ESTIMATE) tool to determine the score of immune and stromal cells [83]. MCPCounter was implemented to calculate the abundance of tissue-infiltrating immune and stromal cell populations in 10 cell types[84]. We used TIDE software with default parameters to analyze the tumor T cell dysfunction and exclusion scoring [85]. To generate a SS specific gene signature matrix in Bulk-RNA seq using immune cell types from the scRNA-seq data, we ran the Create Signature Matrix module (https://cibersortx.stanford.edu/runcibersortx.php) [86] with the log-normalized expression matrix from our dataset restricted to immune populations of interest supplied as a reference matrix. We used a Min. Expression parameter of 1, No. Barcode Genes parameter of 300 to 500, and q-value at 0.01 with all other parameters set to default. Accordingly, CIBERSORTx deconvolution was performed on the SS bulk RNA-seq cohorts in relative mode with S-mode batch correction and quantile normalization disabled to re-impute the cell fractions.

### Non-negative matrix factorization (NMF) clustering

To identify new optimal molecular subtypes of SS, we performed NMF clustering using the R “NMF” package [87] with the parameters as follows: K = 2-10, method = ‘brunet’, run = 50. Log2 transformed TPM data was used as the input data for clustering. Rather than separating clusters based on distance computation, NMF detects context-dependent patterns of gene expression in complex biological systems. To generate the most robust clustering, 25 different truncation points are set from the top 6% gene variance to the top 30% gene variance. We used the cophenetic correlation coefficient to determine the clusters that yield the most robust clusters. The cophenetic correlation coefficient is calculated from the consensus matrix of NMF clusters. From the plot of the resonance correlation coefficient versus k, we chose the top point where the coefficient of conjugate correlation decreased the most. The silhouette width was used to evaluate the performance of clustering, which is defined as the ratio of the average distance of each sample from samples in the same cluster to the minimum distance of samples not in the same cluster.

The optimal numbers of clusters were picked from each set and we calculated the metastasis-free survival (MFS) rate across the clusters in three approaches: i). according to the original subtypes generated by NMF; ii). in case of a group with less than or equal to 3 patients, this was combined with the group with the closet prognostic pattern, named as outliers incorporated; iii). by removing outliers from the MFS analysis. By comparing *p*-value significance (log-rank test, *p* < 0.05) in all three groups, candidates were retained and forwarded to further observe their clustering consensus matrix and overall survival (OS) rate to derive the most robust clusters. The validity of clusters was visualized by Uniform Manifold Approximation and Projection (UMAP) analysis.

### Weighted correlation network analysis (WGCNA) and co-expressed network construction

The R software package “WGCNA” [88] was used to identify the co-expression modules. Genes with the top 25% variance were retained and subjected to clustering analysis. The co-expression network fits a scale-free network with β values ranging from 1 to 20. Also, a linear model was built by the adjacency of the nodes (log k) and the logarithm of the occurrence probability of the nodes [log(p(k))] with a correlation greater than 0.85. The nearest soft threshold was chosen for filtering the co-expression module in order to ensure a scale-free network. Next, the expression matrix was transformed to an adjacency matrix and thereafter to a topological matrix. In accordance with the topological overlap matrix (TOM), we used the mean linkage hierarchical clustering method to keep a minimum number of 30 genes in each module according to the hybrid dynamic cutting tree criterion. Afterwards, we calculated the eigengenes of each module, performed a cluster analysis of the modules, and computed the correlation between modules and subtypes. Hallmark gene sets, Reactome gene sets, and all GO gene sets were employed for functional annotations of the identified modules.

### ScRNA-seq pre-processing

The raw counts of scRNA-seq data were processed with the Seurat package (v 4.10) [89] for each individual sample. The cells with number of expressed genes < 300 and the gene expressed in less than 3 cells were filtered out. In addition, percentages of mitochondrial genes < 15%, ribosome genes > 1%, and hemoglobin <1 % of total expressed genes were retained. In terms of the specific genes, we excluded the housekeeping gene MALAT1 and mitochondrial genes from the downstream analysis.

### ScRNA-seq data analysis

For each sample, gene expression was denoted as a fraction of that gene and multiplied by 10,000, converted to the natural logarithm and normalized after adding 1 to avoid log 0 values. The batch effects were removed by the Harmony package (version 0.1.0) [90] of R. We applied the “FindNeighbors” function in Saruat, using the 1st to 15th harmony dimensions and then looking for clusters at resolutions of 0.01, 0.05, 0.1, 0.2, 0.3, 0.5, 0.8 and 1. We identified clusters with resolution of 0.8 and calculated the UMAP coordinates using the same harmony dimensions for visualization. To annotate cell clusters, we firstly filtered genes to include only those expressed in at least 25% of cells in at least one cluster at a given clustering resolution, and then “FindAllMarkers” function was executed with default parameters to find the cluster markers. Cell groups are annotated according to DEGs and canonical cellular markers from the literature. The cellular markers were listed in Supplementary Table 4.

Single cell pseudotime trajectories were performed using the “Monocle2” package [91] (v2.18.0) in R. The original UMI count-scale gene-cell matrix from the Seurat-processed data was used as input. A newCellDataSet function was applied to create an object with expressionFamily=negbinomial.size and other default parameters. Only genes with mean expression ≥ 0.1 and genes with expression > 10 cells were selected for trajectory analysis. DEGs with q-values <0.01 between cell groups were applied for dimensional reductions using the “reduceDimension” function with parameters reduction method = “DDRTree” and max components = 2. Cells were sorted and visualized with the function “plot_cell_trajectory”.

We used CellChat package (version 1.1.3) [92] to explore the cell-to-cell interactions between malignant cells and immune cells and identify the role of ligands-receptors in specific signaling pathways. The graphical visualization parameter was set at nPatterns = 5.

### DNA sequencing

#### Library preparation and sequencing

A total of 20-250 ng DNA from each sample was used for library preparation with KAPA HyperPlus (Roche) with the following modifications: fragmentation with 12.5 min incubation, xGen Duplex Seq adapters (3-4 nt unique molecular identifiers (UMI), 0.55 µM (for input amounts of 25-250 ng) or 0.15µM (for input amounts of < 25 ng), Integrated DNA Technologies) were used for the ligation, and xGen Indexing primers (2 mM, with unique dual indices, Integrated DNA Technologies) were used for PCR amplification (5-13 cycles depending on input amount of DNA). Target enrichment was performed in a multiplex fashion with a library amount of 375 ng (4-plex). The libraries were hybridized to the capture probe using the Genomic Medicine Sweden mini panel (c389 genes GMS mini / GMCK v1) (Twist Bioscience) with the addition of Twist Universal Blockers and Blocking solution for 16 hours (Supplementary Table 5). The post-capture PCR was performed with xGen Library Amp Primer (0.5 mM, Integrated DNA Technologies) for ten cycles. Quality control was performed with the Quant-iT dsDNA HS assay (Invitrogen) and TapeStation HS D1000 assay (Agilent). Sequencing was done on a NovaSeq 6000 (Illumina) using paired-end 150 nt readout, aiming at 40 M read pairs per sample. Demultiplexing was done using the Illumina bcl2fastq Conversion Software v2.20.

### Panel sequencing data analysis-mutation calling

To identify small tumor variants and single nucleic variants (SNVs), we used the BALSAMIC pipeline v7.2.2[93] to analyze each of the FASTQ files. Firstly, quality control of fastq files was assessed using the FastQC software v 0.11.5[65], after which we trimmed adapter sequences and low-quality bases using the fastp v 0.20.0[94]. Summarized quality results were created by MultiQC v 1.7[95]. The trimmed reads were aligned to the reference genome (Hg19) using BWA MEM v 0.7.15[96]. The resulting SAM files were converted into BAM files and then sorted and indexed with Samtools v 1.6[97]. The duplicate reads were marked using the Picard tools v2.17.0 [98] with MarkDuplicates and eliminated from the downstream analyses and promptly quality controlled using the CollectHsMetrics, CollectInsertSizeMetrics, and CollectAlignmentSummaryMetrics functionalities. For each sample, somatic mutations were called in paired variant calling mode using the VarDict v 2019.06.04[99] and annotated using the Ensembl VEP v 99.1[100]. For tumor mutational calling, all low-quality variants were initially removed via a series of variant filtering using bcftools v 1.9.0[101], including Mean Mapping Quality (MQ) >= 40. Total read depth (DP) >= 100, Variant Depth (VD) >=5.0, Allele frequency (AF) >= 0.01, and Maximum allele frequency across populations < 0.005. All passed variants were converted to Mutation Annotation Format (MAF) format using vcf2maf, with the annotation added by Ensembl VEP v 99.1 [102]. Subsequently, additional filtering was performed to reduce false-positive calls and keep the somatic mutations: 1) variant allele frequency (VAF) > 10%; 2) Somatic != NULL; 3) Impact Variants = Moderate or high. Variants classified as putative functionally relevant and as of unknown/contradictory functional significance were selected, whereas neutral variants, alternations without representative gene/transcript, and polymorphisms were excluded from the downstream analyses by using the Molecular Tumor Board (MTB) Portal[103] (accessed 11/2021). The latter is a clinical decision support tool that provides a general interpretation of a given list of cancer gene variants that combines the latest results from clinical and preclinical studies, true biological hypotheses, and bioinformatics calculations to classify a variant as biologically relevant and to evaluate the functional and predictive relevance of genomic alterations. The TMB of each tumor was calculated with the “tmb” function of the R package “maftools” [63].

### Somatic Copy Number Alterations (SCNAs)

CNVkit v 0.9.4a0[104] was used with default parameters on paired tumor-normal sequencing data. The log2 copy ratio in the cnr.file from the CNVkit program was adjusted to represent the length of the bin, and copy number segments from the given coverage table using default function circular binary segmentation (CBS). Focal and broad copy number variations were detected using the CNVkit segmentation data imported in GISTIC 2.0[105] with the following parameters. (-ta 0.2 -td 0.25 - genegistic 1 - maxseg 3200 -broad 1 -conf 0.99 -rx 0 –brlen 0.7 -cap 1.5 –armpeel 1).

For the differential region analysis, we used a two-step procedure inspired by the Korthauer and the Shih methods for detecting regions of differential methylation and SCNA regions[106, 107]. Since q-values were calculated from the results of two paired sets of GISTICs, they were divided into two groups: i) overlapped significant GISTIC cytobands, which took the smaller q-value result; ii) for non-overlapping, the original GISTIC results were searched in another set, and if a GISTIC score appeared, it is added and the q-value is prevailed over the significant non-overlapping result. In terms of the false-positive results generated by GISTIC (where one cytoband may correspond to multipal chromosome regions, and genes may match in the wrong locations), to capture the significant drivers, genes were placed according to their physical location on the chromosome band. The genome annotation (hg19/GRCh37) obtained from the UCSC database (http://hgdownload.cse.ucsc.edu/goldenpath/hg19/database/), and BEDTools [108] were then applied to map the locations of differential regions and genes screened out as true positives.

### HiRIEF-nanoLC-MS/MS based proteomics

A total of 14 fresh frozen SS samples were dissolved in 200 μl lysis buffer (4% SDS, 50mM Hepes, 1mM DTT, pH 7.4) and the total protein amount was estimated (DC kit, BioRad). Samples were heated and sonicated then prepared for mass spectrometry analysis using a modified version of the SP3 protein clean-up and a digestion protocol [109], where proteins were digested by LysC and trypsin (sequencing grade modified, Pierce). In brief, around 100 µg protein from each sample was alkylated with 4 mM chloroacetamide, sera-Mag SP3 bead mix (20 µl) was added to the protein sample together with 100 % acetonitrile to a final concentration of 60 %, and the mix was incubated under rotation at room temperature for 20 min. The mix was then placed on a magnetic rack and the supernatant was discarded, followed by two washes with 70 % ethanol and one with 100 % acetonitrile. The beads-protein mixture was reconstituted in 100 μl LysC buffer (0.5 M Urea, 50 mM HEPES pH: 7.6 and 1:50 enzyme (LysC) to protein ratio) and incubated overnight. Finally, trypsin was added in 1:50 enzyme to protein ratio in 100 μl 50 mM HEPES pH 7.6 and incubated overnight. The peptides were eluted from the mixture after placing the mixture on a magnetic rack, followed by peptide concentration measurement (DC kit, BioRad). The samples were then pH adjusted using TEAB pH 8.5 (100 mM final conc.), 40 μg of peptides from each sample were labelled with isobaric TMT-tags (TMTpro 16 plex reagent) according to the manufacturer’s protocol (Thermo Scientific), and 320 μg of peptides separated by immobilized pH gradient - isoelectric focusing (IPG-IEF) on 3-10 strip and 3.7-4.9 strip as described previously [110].

Labelling efficiency was determined by LC-MS/MS before pooling of the samples. For the sample clean-up step, a solid phase extraction (SPE strata-X-C, Phenomenex) of TMT labelled and pooled peptides was performed and purified samples were dried in a SpeedVac. An aliquot of approximately 10 μg was suspended in LC mobile phase A and 2 μg was injected on the LC-MS/MS system. Online LC-MS was performed as previously described [110] using a Dionex UltiMate™ 3000 RSLCnano System coupled to a Q-Exactive-HF mass spectrometer (Thermo Scientific). Each of the 72 fractions in the plate wells was dissolved in 20 μL solvent A and 10 μL were injected. Samples were trapped on a C18 guard-desalting column (Acclaim PepMap 100, 75μm x 2 cm, nanoViper, C18, 5 μm, 100 Å), and separated on a 50 cm long C18 column (Easy spray PepMap RSLC, C18, 2 μm, 100 Å, 75 μm x 50 cm). The nano capillary solvent A was 95 % water, 5% DMSO, 0.1 % formic acid; and solvent B was 5 % water, 5 % DMSO, 95 % acetonitrile, 0.1 % formic acid. At a constant flow of 0.25 μl min−1, the curved gradient went from 6-10 % B up to 40 % B in each fraction in a dynamic range of gradient length, followed by a steep increase to 100 % B in 5 min. FTMS master scans with 60,000 resolution (and mass range 300-1500 m/z) were followed by data-dependent MS/MS (30 000 resolution) on the top 5 ions using higher energy collision dissociation (HCD) at 30 % normalized collision energy. Precursors were isolated with a 2 m/z window. Automatic gain control (AGC) targets were 1e6 for MS1 and 1e5 for MS2. Maximum injection times were 100 ms for MS1 and 100 ms for MS2. The entire duty cycle lasted ∼2.5 s. Dynamic exclusion was used with 30 s duration. Precursors with unassigned charge state or charge state 1 were excluded. An underfill ratio of 1 % was used.

Orbitrap raw MS/MS files were converted to mzML format using msConvert from the ProteoWizard tool suite [111]. Spectra were then searched using MSGF+ (v10072) [112] and Percolator (v2.08) [113]. All searches were done against the human protein subset of Ensembl (ENS104) in the Nextflow platform (https://github.com/lehtiolab/ddamsproteomics, vs1.5).

Pairwise subtype differences at the protein level were estimated with DEqMS (v1.12.1)[114]. Enrichment analysis was performed on the sorted log2 fold changes with fgsea (v1.20.0) [115] using MSigDB Reactome canonical pathways v7.4 [79] and significant pathways were called at an FDR < 0.01.

### OpalTM multiplexing and digital image analysis

OpalTM Multiplexing reagents (Akoya Biosciences) were utilized to detect potential co-expression of multiple markers in the same cell in situ. Formalin-fixed, paraffin-embedded 4 μm tumor sections from 21 patients were utilized for multiplexing staining. Deparaffinization and rehydration (Decloaking NxGen Chamber BioCare Medical) were performed before the heat-induced epitope retrieval (Citrate Buffer pH 6.0, C9999, Sigma-Aldrich) at 110°C for 5 minutes. A 10-minute incubation blocking step (Blocking diluent, ARD1001EA, Akoya Biosciences) was performed followed by primary antibody incubation. Primary antibodies included: CD20 (M0755, Dako), CD4(ab133616), CD3(85061, Cell Signaling), CD8(M7103, Dako), and PD1(86163, Cell Signaling). Secondary antibody (Opal Polymer anti-Ms and Rb HRP, Akoya Biosciences) was applied for incubation for 10 minutes at room temperature. Opal fluorophores were diluted in amplification diluent (FP1498, Akoya Biosciences) at 1:150 and followed by an incubation of 10 minutes at room temperature avoiding light. Between each round of antibody and after the last round of staining, a heat-induced epitope retrieval (Citrate Buffer pH 6.0, C9999, Sigma-Aldrich) was performed for 20 min at 95°C. Slides were mounted with ProLongTM Glass Antifade Mountant with NucBlueTM stain (P36985, ThermoFisher).

Imaging was performed with the Vectra Polaris scanning system at 40x magnification. Five random areas with a size of 2000*1500 μm were captured for analysis in each sample. The Qupath software (0.3.2) was used for cell segmentation and classification. Cell detection was performed using the default settings in Qupath: Threshold: 2, requested pixel size: 0.227 μm, background radius: 8 μm, median filter :0 μm, minimum area:10,000 μm2, maximum area: 400,000 μm2 and cell expansion: 2 μm. Subsequently, a single classifier for each channel was set up by manual adjustment according to the training area and a combined classifier was established. The combined classifier was applied to all images and the results were exported into a .txt file for statistical analysis.

### Statistics

The unpaired Student’s t-test was used to analyze the comparison between two continuous variables and a normally distributed variable. Non-normally distributed variables were analyzed with the Mann-Whitney U test. The Kaplan-Meier method in the R package “survminer” and “survival” was used for survival analysis with the log-rank test for comparisons between groups. In order to compare three or more groups, ANOVA and Kruskal-Wallis tests were performed on parametric and non-parametric variables respectively. Correlations of rank-ordered values were assessed using the Spearman rank correlation test. Statistical analysis was conducted in the R version 4.0.3 (R Foundation for Statistical Computing, Vienna, Austria).

## Extended Data

**Extended Data Figure. 1:**
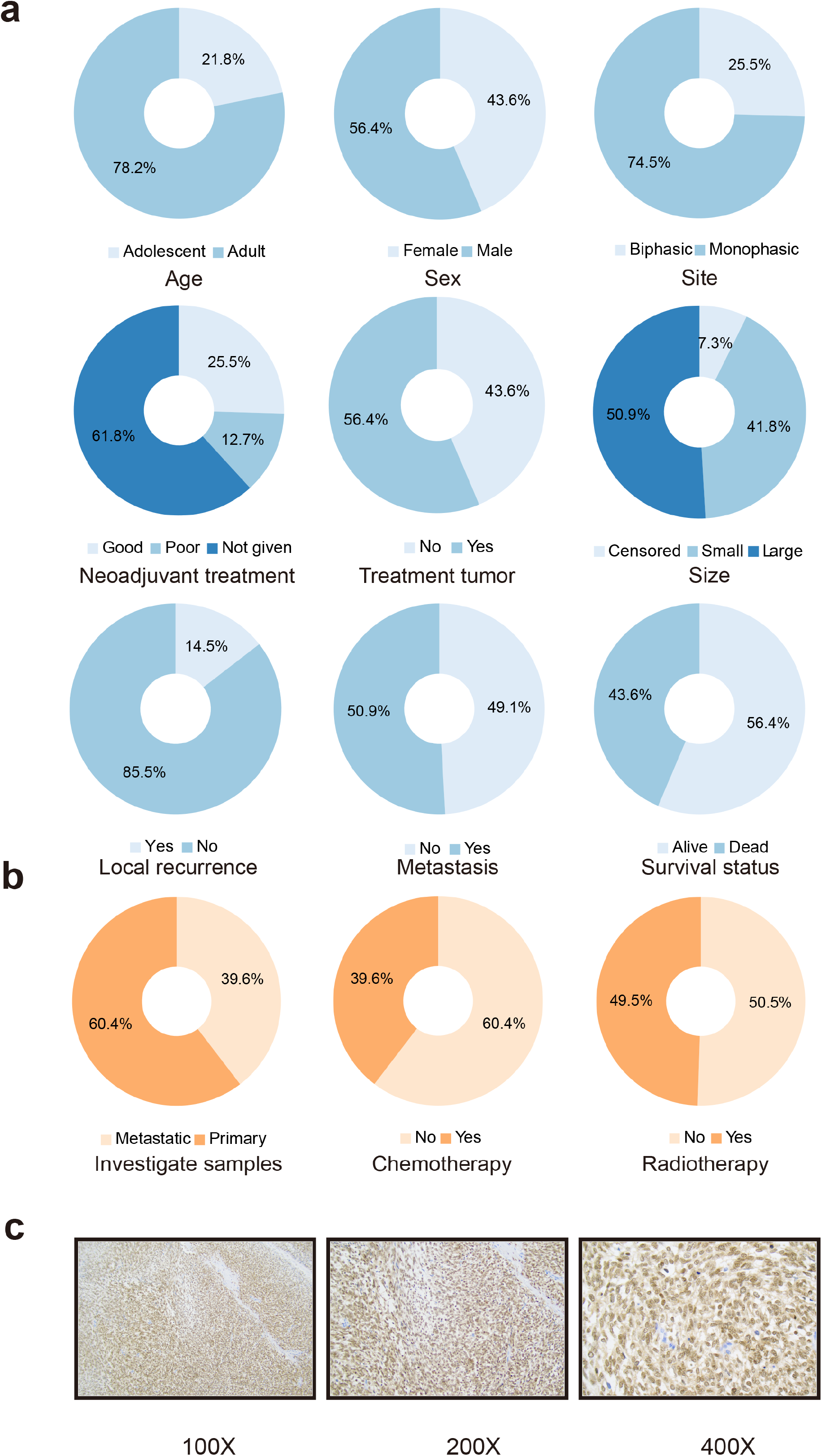
**(a-b).** Summary of clinical characteristics of **(a)** 55 synovial sarcoma patients and **(b)** 91 patients. **(c)** SS18-SSX immunoreactivity at 100 ×, 200 ×, and 400 × magnifications.

**Extended Data Figure. 2:**
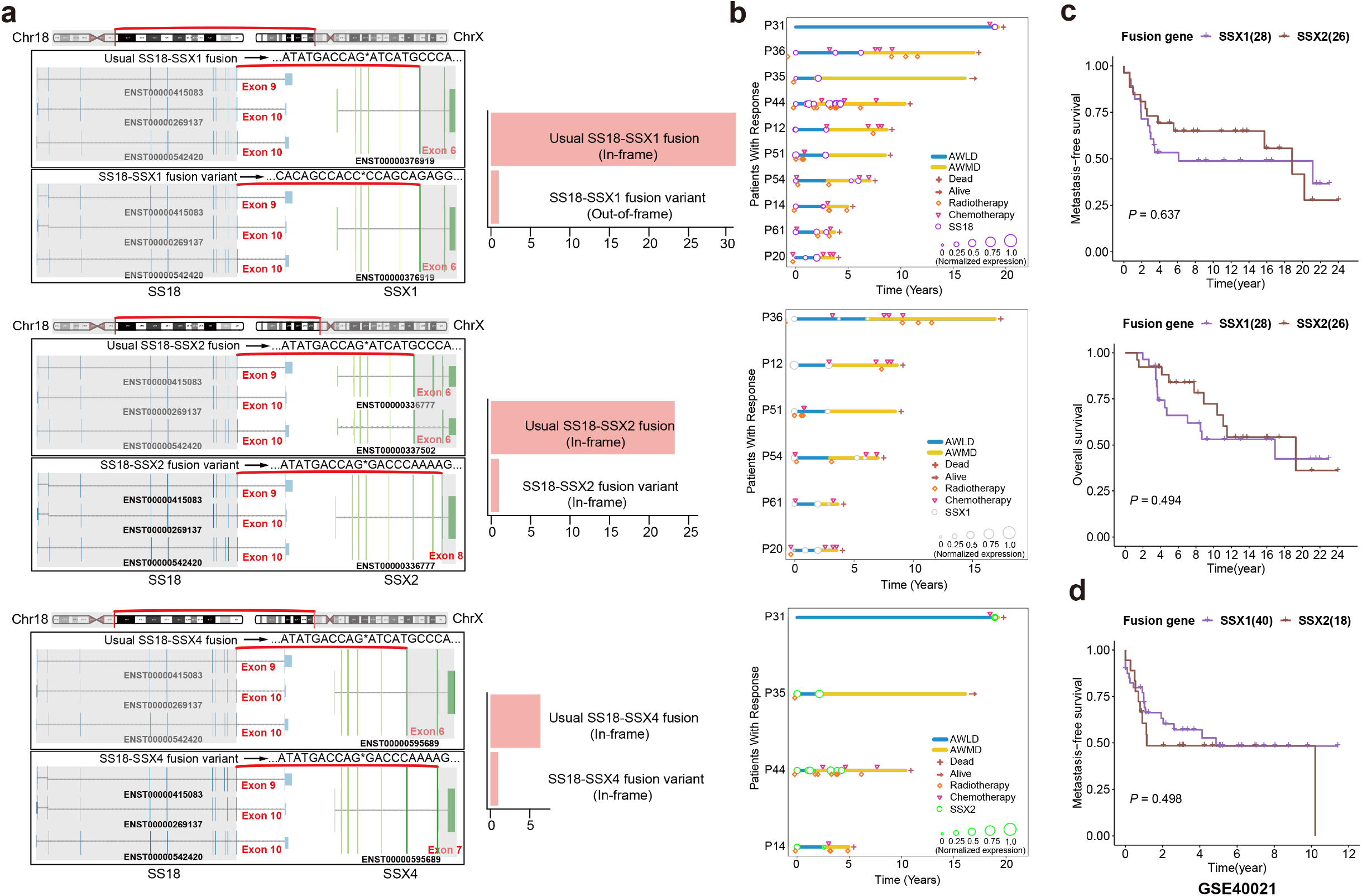
**(a)** The landscape of SS18-SSX (1,2,4) variants in the transcripts level and their proportions. **(b).** Assessment of changes in the characteristics of SS18, SSX1, and SSX2 expressions with treatment response in longitudinal analysis. Blue and yellow bars indicate the primary and metastatic status, and the size of circles represents the expression levels, respectively. AWLD, alive with local disease, AWMD, alive with metastatic disease.

**Extended Data Figure. 3:**
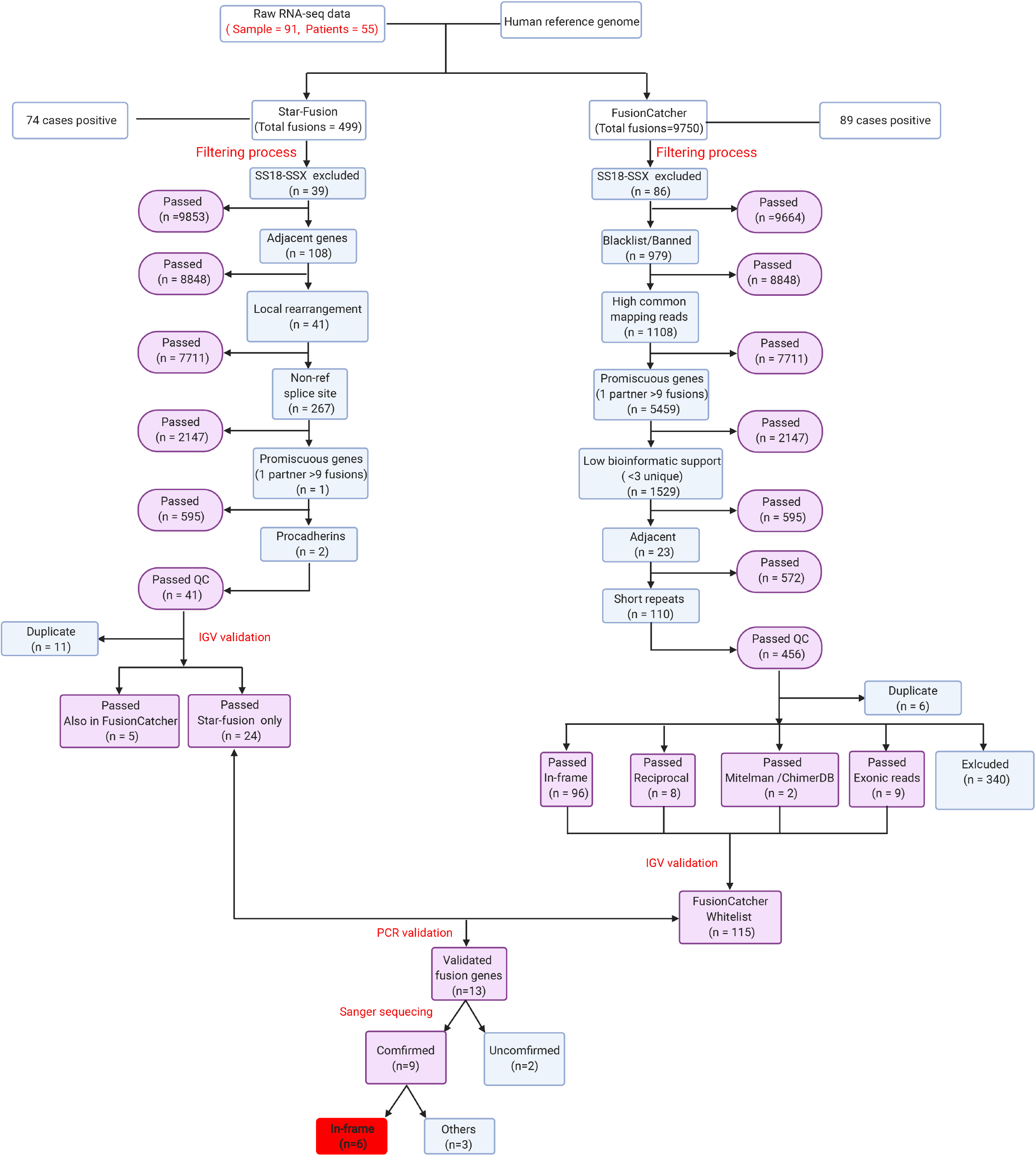
Workflow of the fusion gene filtering process.

**Extended Data Figure. 4:**
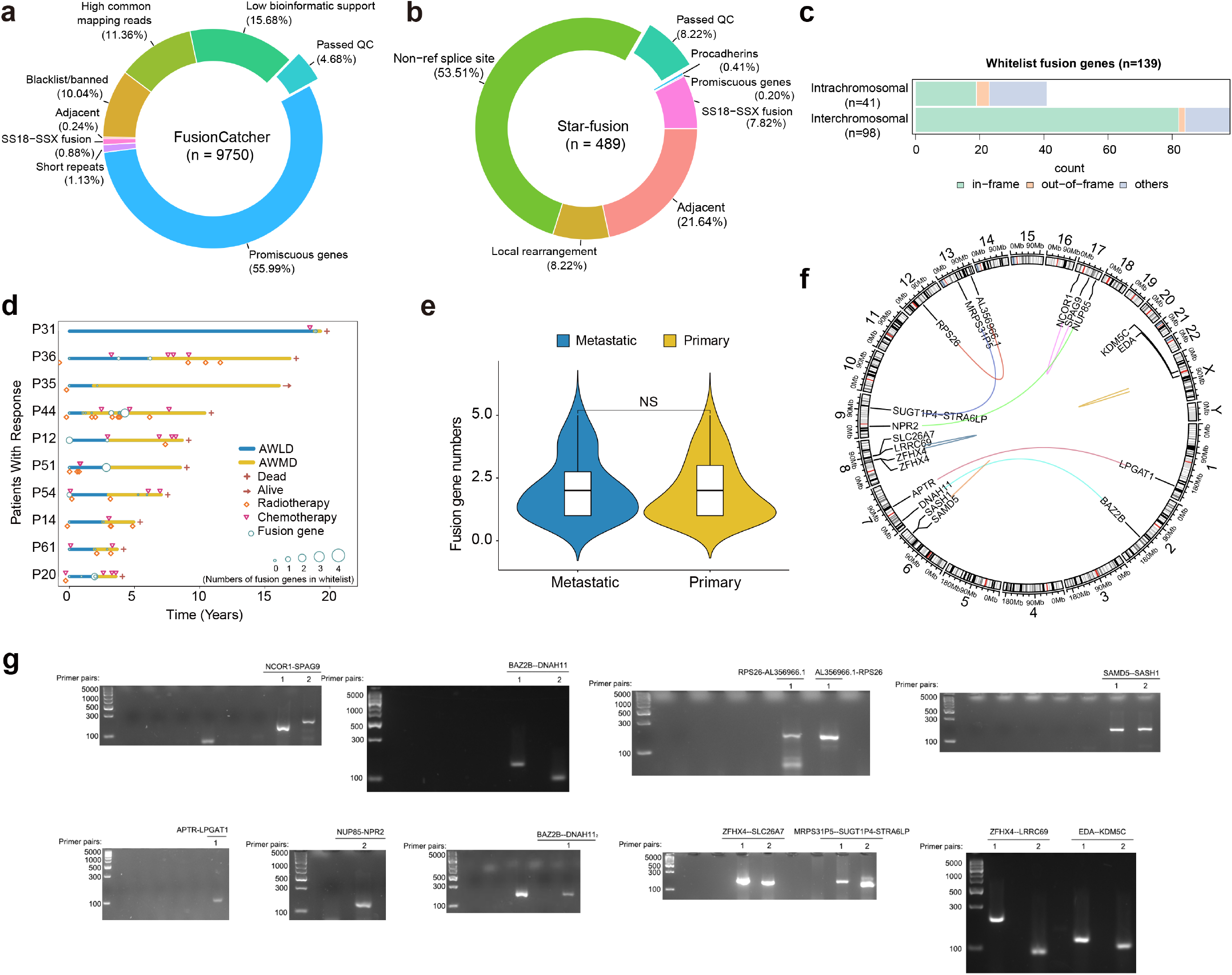
**(a-b).** Proportions of fusion gene types generated by **(a)** FusionCatcher and **(b)** Star-fusion. **(c).** Types of the fusion gene in the whitelist. **(d).** Assessment of changes in the characteristics of whitelist fusion gene numbers with treatment response in longitude analysis. Blue and yellow bars indicate the primary and metastatic status, the size of circles represents the expression levels, respectively. **(e).** Distributions of whitelist fusion gene numbers between metastatic and primary samples. Middle line: median; box edges: 25th and 75th percentiles. Mann-Whitney U test. **(f).** Circos plot depicting the validated fusion genes. **(g).** PCR validated whitelist fusion genes.

**Extended Data Figure. 5:**
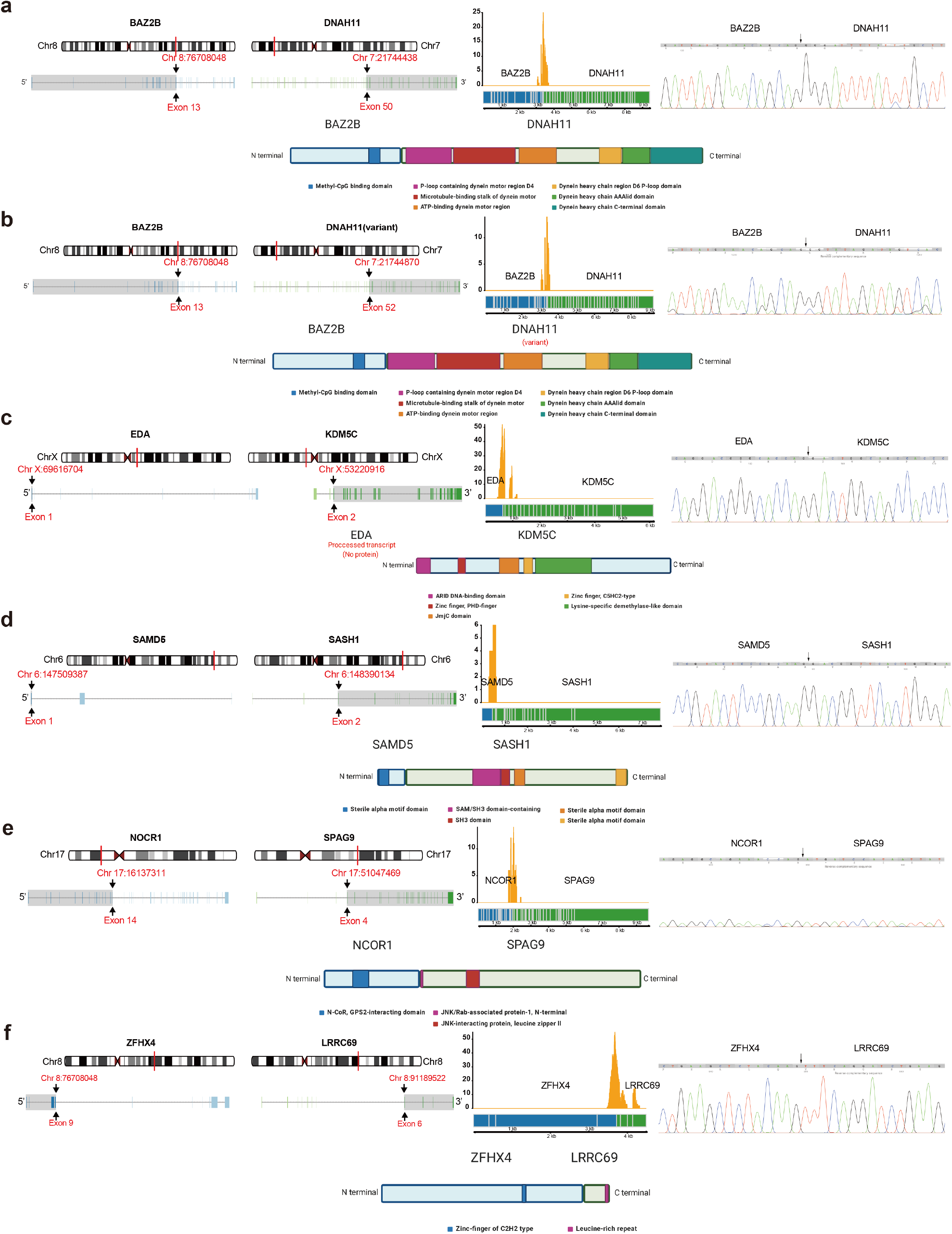
**(a-f).** True positive secondary fusion genes were confirmed by sanger sequencing.

**Extended Data Figure. 6:**
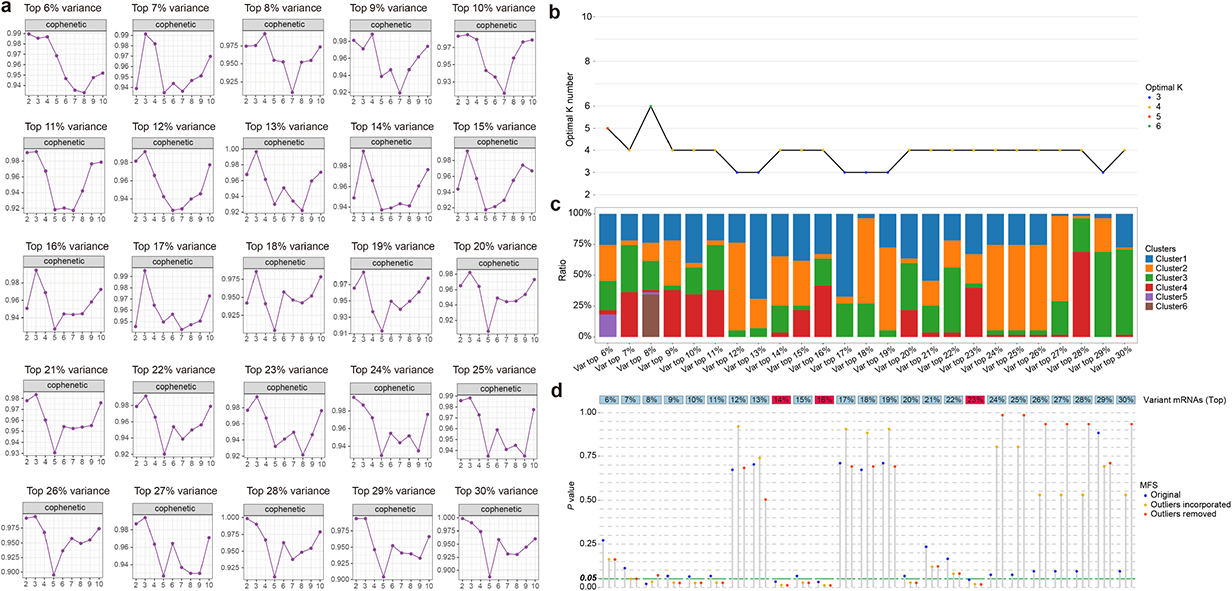
**(a).** Cophenetic correlation plots from top 6% to top 30% variance in mRNAs. **(b).** NMF clustering of the distribution of the optimal K numbers from the top 6% to the top 30% variant mRNAs. **(c).** NMF clustering of the distribution of the number of patients from the top 6% to the top 30% variant mRNAs. **(d).** NMF clustering of the distribution of the MFS values (original, outliers incorporated, outliers removed) from the top 6% to the top 30% variant mRNAs.

**Extended Data Figure. 7:**
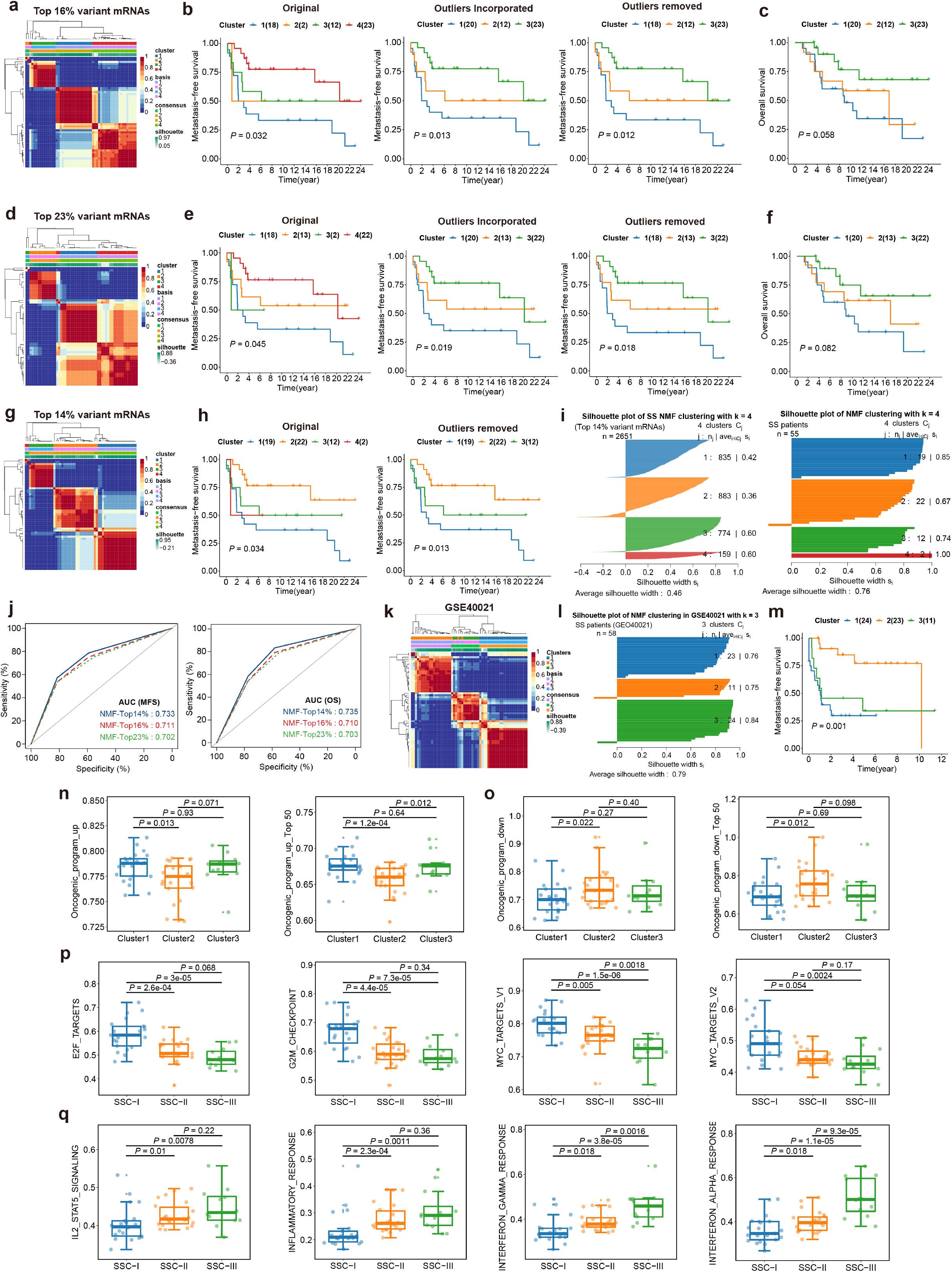
**(A).** Consensus matrix of NMF clustering at top 16% mRNAs variance. **(B).** Kaplan-Meier curve of clusters of metastasis-free survival in synovial sarcoma patients (original, outliers incorporated, outliers removed) at top 16% mRNAs variance (Log-rank test, *P* = 0.014, 0.013, and 0.012 respectively). **(C).** Kaplan-Meier curve of overall survival in synovial sarcoma patients with clusters outliers incorporated at top 16% mRNAs variance (Log-rank test, *P* = 0.058). **(D).** Consensus matrix of NMF clustering at top 23% mRNAs variance. **(E).** Kaplan-Meier curve of clusters of metastasis-free survival in synovial sarcoma patients (original, outliers incorporated, outliers removed) at top 23% mRNAs variance (Log-rank test, *P* = 0.045, 0.019, and 0.018 respectively). **(F).** Kaplan-Meier curve of overall survival in synovial sarcoma patients with clusters outliers incorporated at top 23% mRNAs variance (Log-rank test, *P* = 0.082). **(g-h).** Kaplan-Meier curve of clusters of metastasis-free survival in synovial sarcoma patients (**g)** original, **(h)** outliers removed) at top 14% mRNAs variance (Log-rank test, *P* = 0.034, and 0.013 respectively). **(i).** Silhouette plot of NMF clustering of genes and patients at top 14% mRNAs variance. **(j).** ROC curves of the predication performances of top 14%, 16%, and 23% variances in MFS and OS, respectively. **(k).** Consensus matrix of NMF clustering at top 14% mRNAs variance applied at GSE40021 validation cohort. **(l).** Silhouette plot of NMF clustering of patients at top 14% mRNAs variance applied at GSE40021 validation cohort. **(m).** Kaplan-Meier curve of clusters of metastasis-free survival in patients at top 14% gene variance applied at GSE40021 validation cohort (Log-rank test, *P* = 0.01). **(n-o).** Distributions of the ssGSEA score of the *core oncogenic program* **(n)** upregulated genes and **(o)** downregulated genes in the three SSCs at GSE40021. Middle line: median; box edges: 25th and 75th percentiles. Mann-Whitney U test. **(p-q).** Distributions of the ssGSEA score of **(p)** four significant proliferative and **(q)** immune-related hallmarks in the three SSCs. Middle line: median; box edges: 25th and 75th percentiles. Mann-Whitney U test.

**Extended Data Figure. 8:**
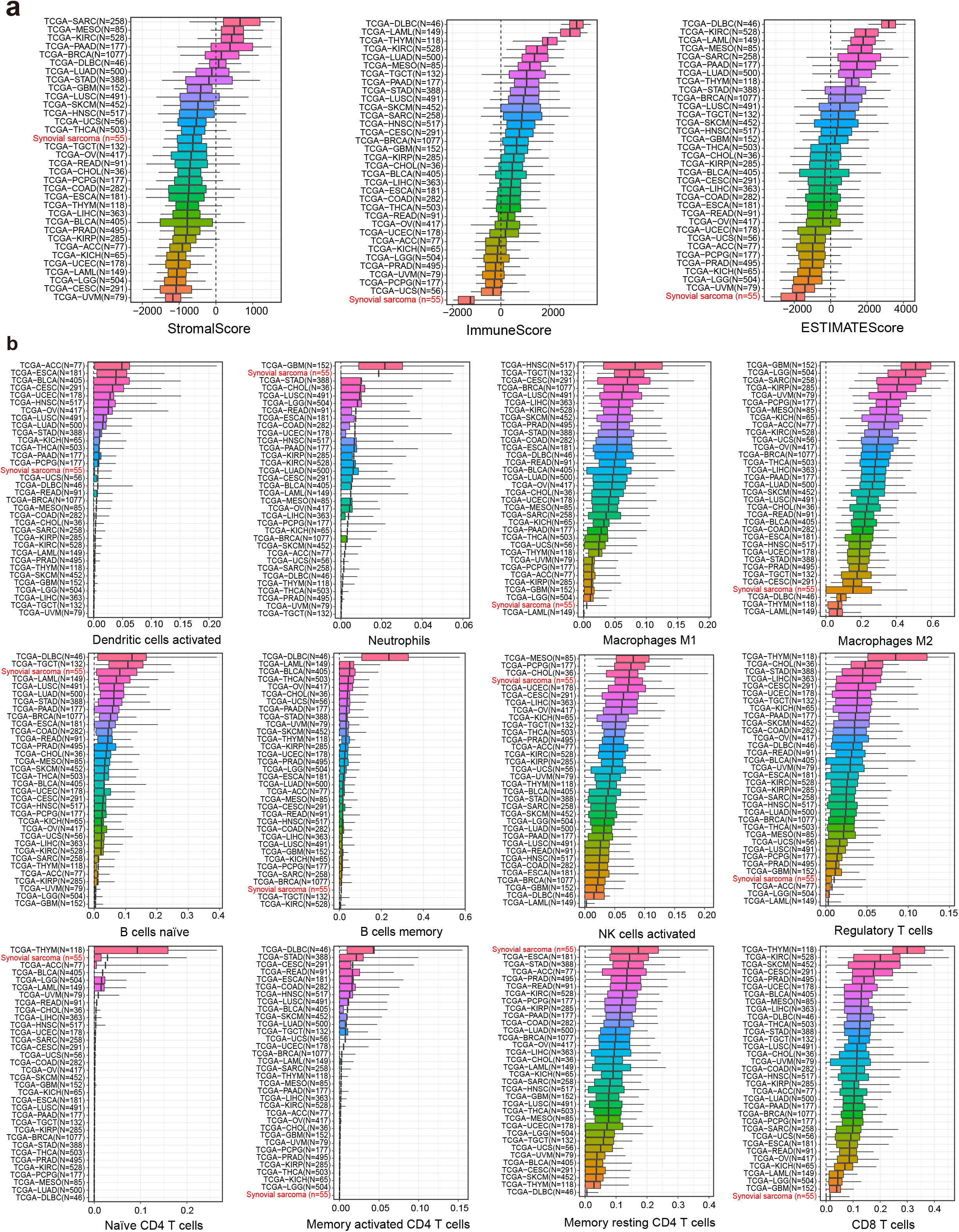
**(a).** Distributions of stromal and immune infiltration abundance by ESTIMATE algorithm of 55 synovial sarcomas and 33 cancer types from TCGA data. **(b).** Distributions of immune cell abundance by CIBERSORT algorithm of 55 synovial sarcomas and 33 cancer types from TCGA data. Middle line: median; box edges: 25th and 75th percentiles. ACC, adrenocortical carcinoma; BLCA, bladder Urothelial Carcinoma; BRCA, breast invasive carcinoma; CESC, cervical squamous cell carcinoma and endocervical adenocarcinoma; CHOL, cholangiocarcinoma; COAD, colon adenocarcinoma; DLBC, lymphoid neoplasm diffuse large B-cell Lymphoma; ESCA, esophageal carcinoma; GBM, glioblastoma multiforme; HNSC, head and neck squamous cell carcinoma; KICH, kidney chromophobe; KIRC, kidney renal clear cell carcinoma; KIRP, kidney renal papillary cell carcinoma; LAML, acute myeloid leukemia; LGG, brain lower grade glioma; LIHC, liver hepatocellular carcinoma; LUAD, lung adenocarcinoma; LUSC, lung squamous cell carcinoma; MESO, mesothelioma; OV, ovarian serous cystadenocarcinoma; PAAD, pancreatic adenocarcinoma; PCPG, pheochromocytoma and paraganglioma, PRAD, prostate adenocarcinoma; READ, rectum adenocarcinoma; SARC, sarcoma; STAD, stomach adenocarcinoma; SKCM, skin cutaneous melanoma; TGCT, Testicular germ cell tumors; THCA, thyroid carcinoma; THYM, thymoma; UCEC, uterine corpus endometrial carcinoma; UCS, uterine carcinosarcoma; UVM, uveal melanoma.

**Extended Data Figure. 9:**
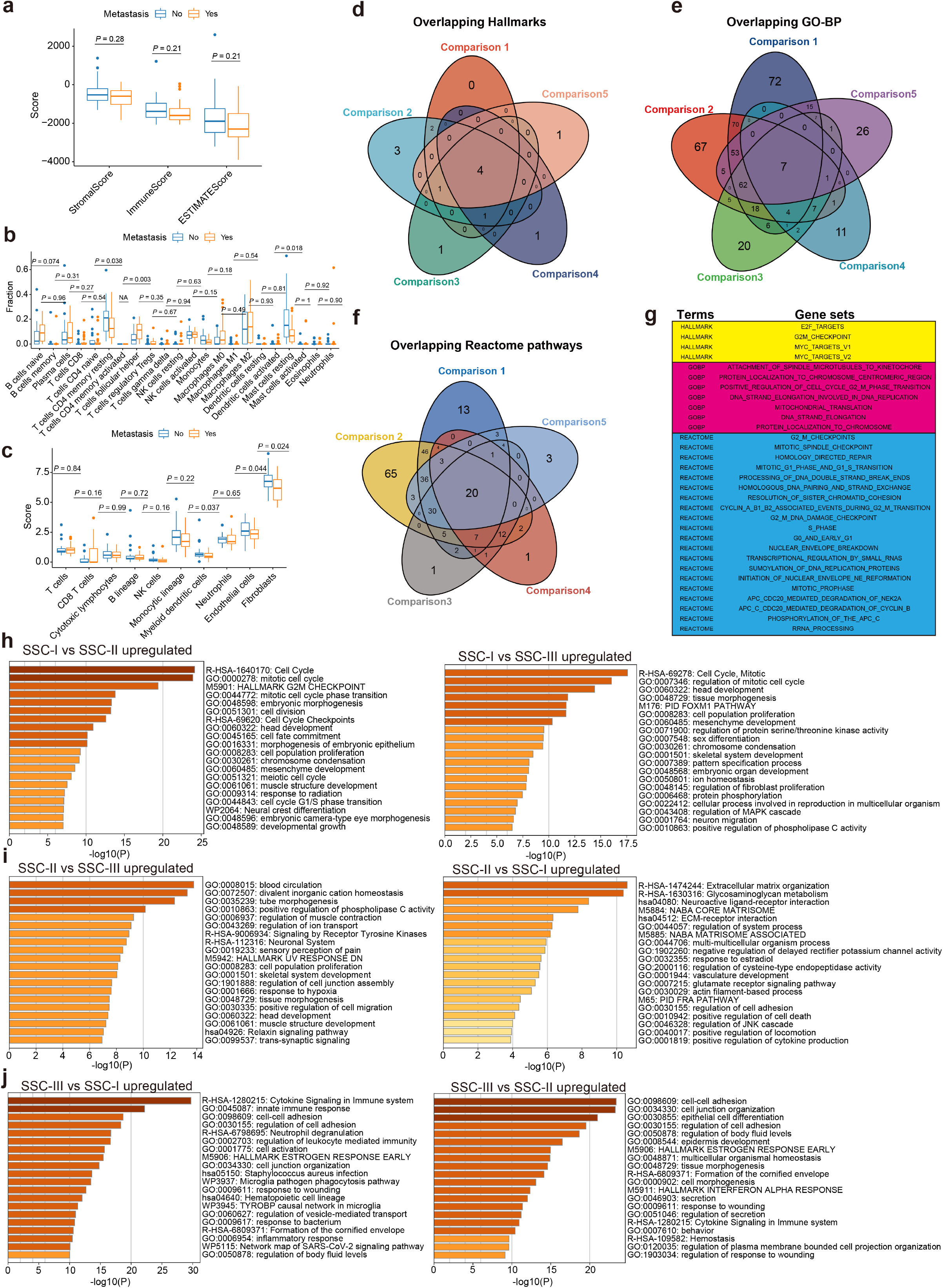
**(a-c).** The distributions of immune and stroma cell types between metastatic and primary patients by three deconvolutional approaches, including **(a)** ESTIMATE, **(b)** CIBERSORT, and **(c)** MCPCounter, respectively, Middle line: median; box edges: 25th and 75th percentiles. Mann-Whitney U test. **(d-f).** Venn diagram depicting the overlap of the overlapping **(d)** Hallmarks, **(e)** GO biological process, and **(f)** Reactome pathways among five comparisons. **(g).** The overlapping gene sets. **(h)**. Functional enrichments of six comparisons of DEGs.

**Extended Data Figure. 10:**
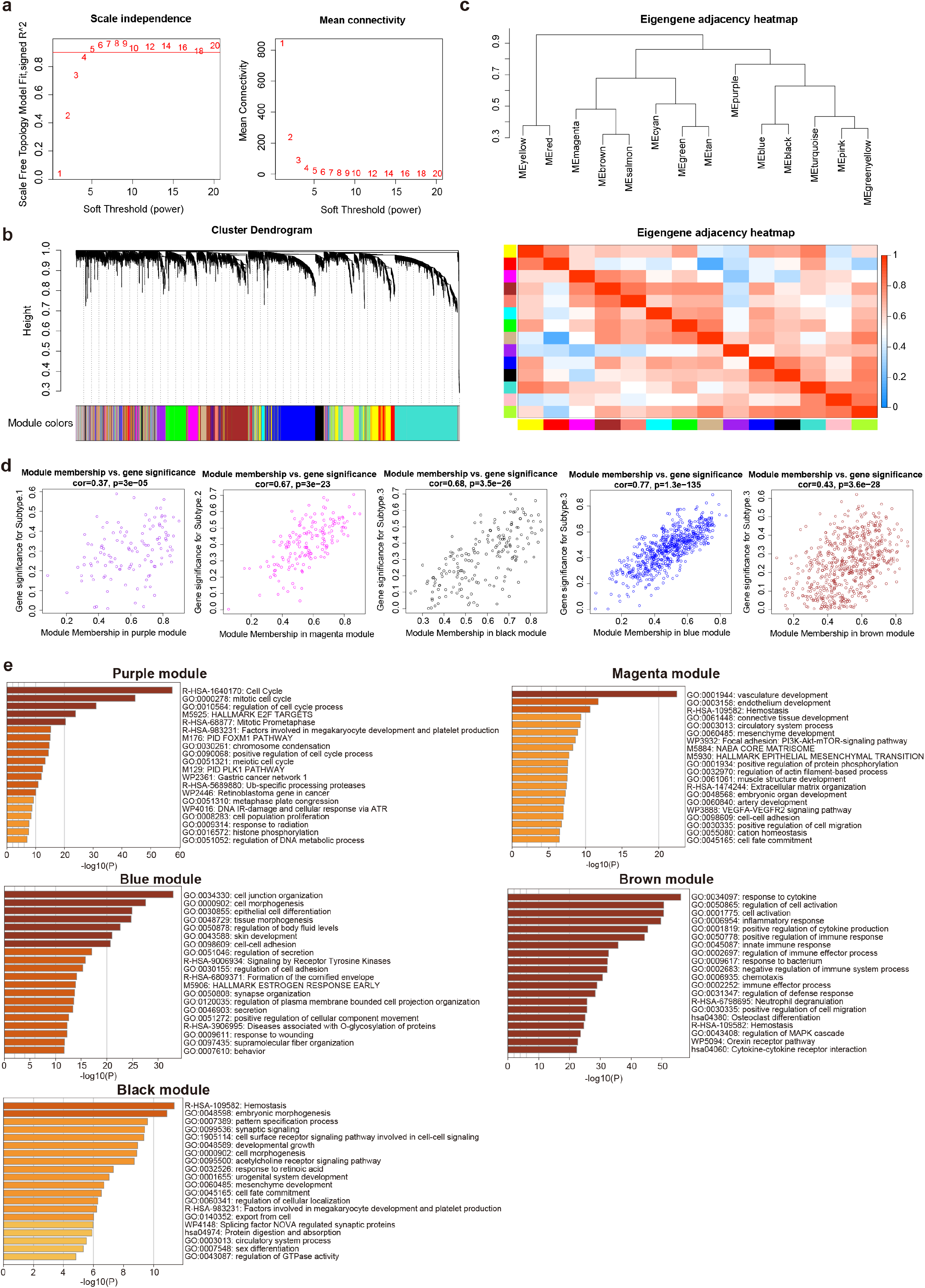
**(a).** Analysis of the scale-free fit index for various soft-thresholding powers (β). **(b).** Dendrogram of all differentially expressed gene modules clustered based on a dissimilarity measure (1-TOM). **(c).** Adjacency heatmap of the gene modules. **(d).** Scatter plot of module members versus gene significance for SSC-I in the purple module; SSC-II in the magenta module; SSC-III in the black, blue, and brown modules. **(e)**. Functional enrichment of five gene modules.

**Extended Data Figure. 11:**
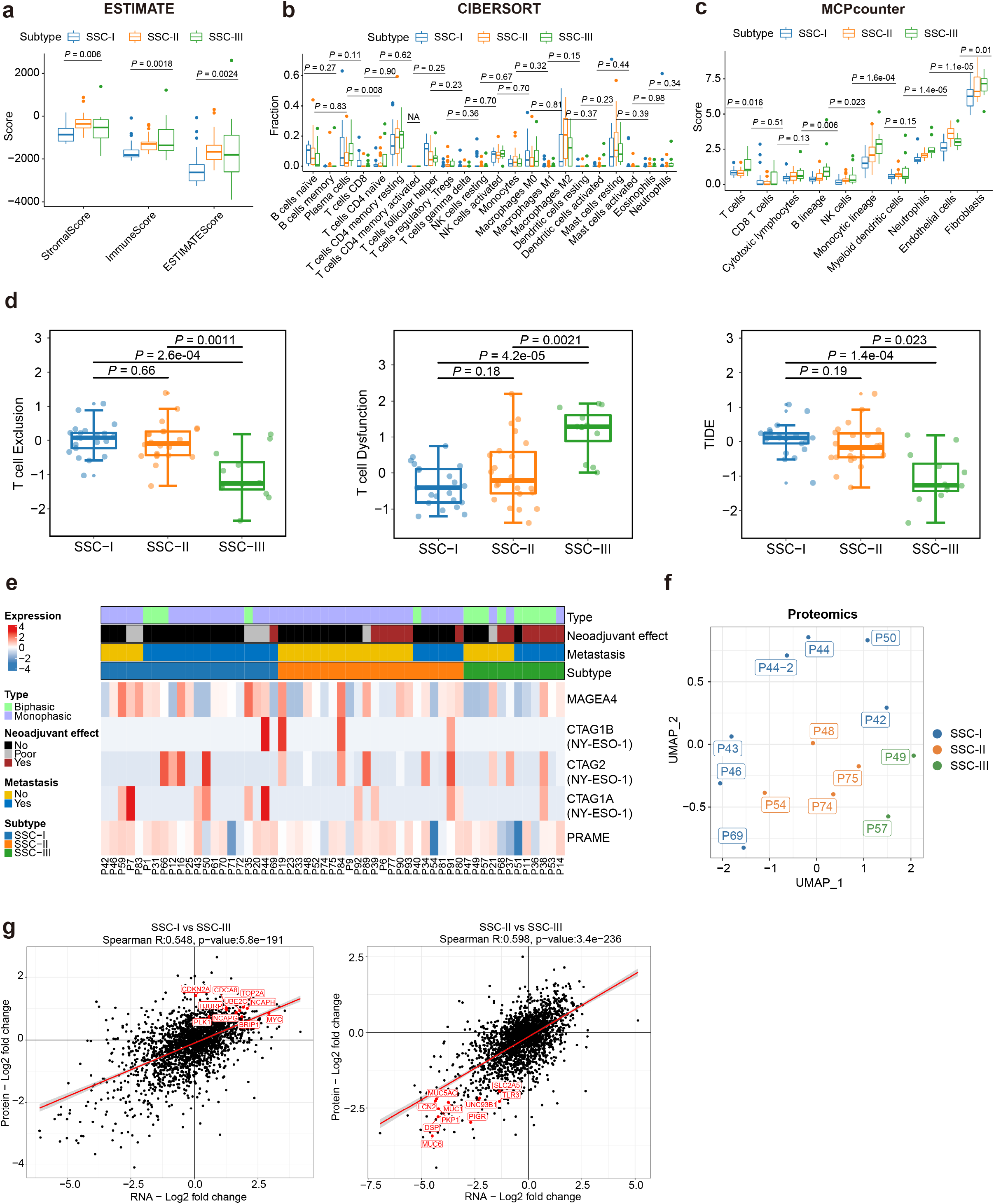
**(a-c).** The distributions of immune and stroma cell types across SSCs by three deconvolutional approaches, including **(a)** ESTIMATE, **(b)** CIBERSORT, and **(c)** MCPCounter, respectively, Middle line: median; box edges: 25th and 75th percentiles. Kruskal-Wallis test. **(d).** Distributions of the T cell dysfunction, T cell exclusion, and TIDE score in three SSCs. Middle line: median; box edges: 25th and 75th percentiles. Mann-Whitney U test. **(e)**. Heatmap of the CTAs in SS patients. Clinicopathological characteristics (top) of the 55 synovial sarcoma patients are shown in the annotation, and different colors represent the characteristics and subtypes. Gene expression (bottom) from 55 patients profiled with CTAs are depicted. **(f).** UMAP plot of SS subtypes in the proteome level. **(g).** The correlation of DEGs between RNA and protein levels, Spearman rank correlation.

**Extended Data Figure. 12:**
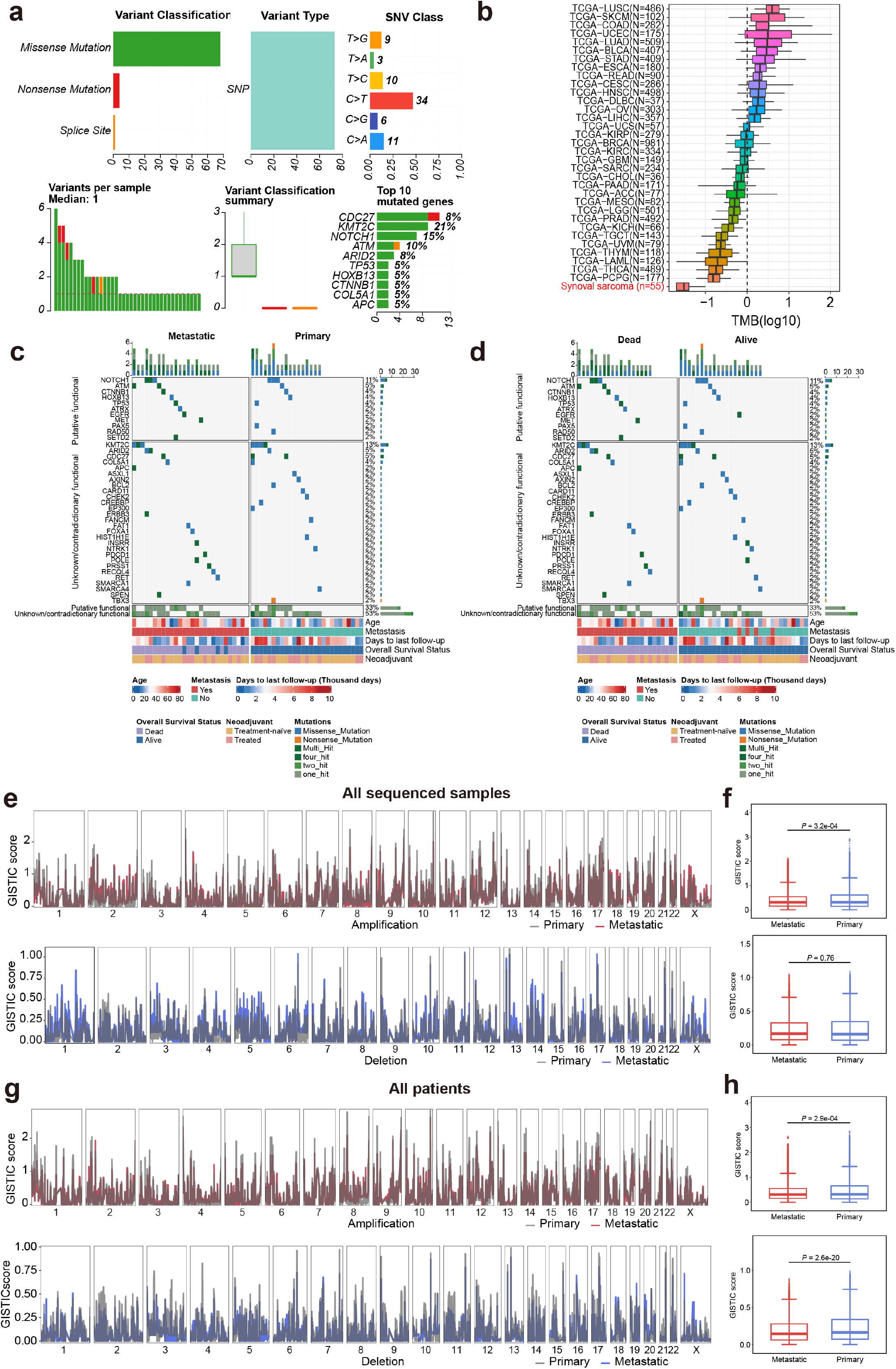
**(a).** Summary of somatic mutation profile in 53 synovial sarcoma patients. **(b).** Distributions of tumor mutation burdens of 53 synovial sarcomas and 33 cancer types from TCGA data. Middle line: median; box edges: 25th and 75th percentiles. ACC, adrenocortical carcinoma; BLCA, bladder Urothelial Carcinoma; BRCA, breast invasive carcinoma; CESC, cervical squamous cell carcinoma and endocervical adenocarcinoma; CHOL, cholangiocarcinoma; COAD, colon adenocarcinoma; DLBC, lymphoid neoplasm diffuse large B-cell Lymphoma; ESCA, esophageal carcinoma; GBM, glioblastoma multiforme; HNSC, head and neck squamous cell carcinoma; KICH, kidney chromophobe; KIRC, kidney renal clear cell carcinoma; KIRP, kidney renal papillary cell carcinoma; LAML, acute myeloid leukemia; LGG, brain lower grade glioma; LIHC, liver hepatocellular carcinoma; LUAD, lung adenocarcinoma; LUSC, lung squamous cell carcinoma; MESO, mesothelioma; OV, ovarian serous cystadenocarcinoma; PAAD, pancreatic adenocarcinoma; PCPG, pheochromocytoma and paraganglioma, PRAD, prostate adenocarcinoma; READ, rectum adenocarcinoma; SARC, sarcoma; STAD, stomach adenocarcinoma; SKCM, skin cutaneous melanoma; TGCT, Testicular germ cell tumors; THCA, thyroid carcinoma; THYM, thymoma; UCEC, uterine corpus endometrial carcinoma; UCS, uterine carcinosarcoma; UVM, uveal melanoma. **(c-d).** Fifty-five synovial sarcoma patients with/without mutation data are ordered by their mutation frequencies and separated by **(c)** metastasis and **(d)** survival status. **(e).** GISTIC amplification (top) and deletion (bottom) plots of primary and metastasis in all sequenced samples. **(f).** GISTIC scores distributions of amplifications and deletions between primary and metastatic in all sequenced samples. Middle line: median; box edges: 25th and 75th percentiles. Mann-Whitney U test. **(g).** GISTIC amplification (top) and deletion (bottom) plots of primary and metastatic in all patients. **(h).** GISTIC scores distributions of amplifications and deletions between primary and metastatic in all patients. Middle line: median; box edges: 25th and 75th percentiles. Mann-Whitney U test.

**Extended Data Figure. 13:**
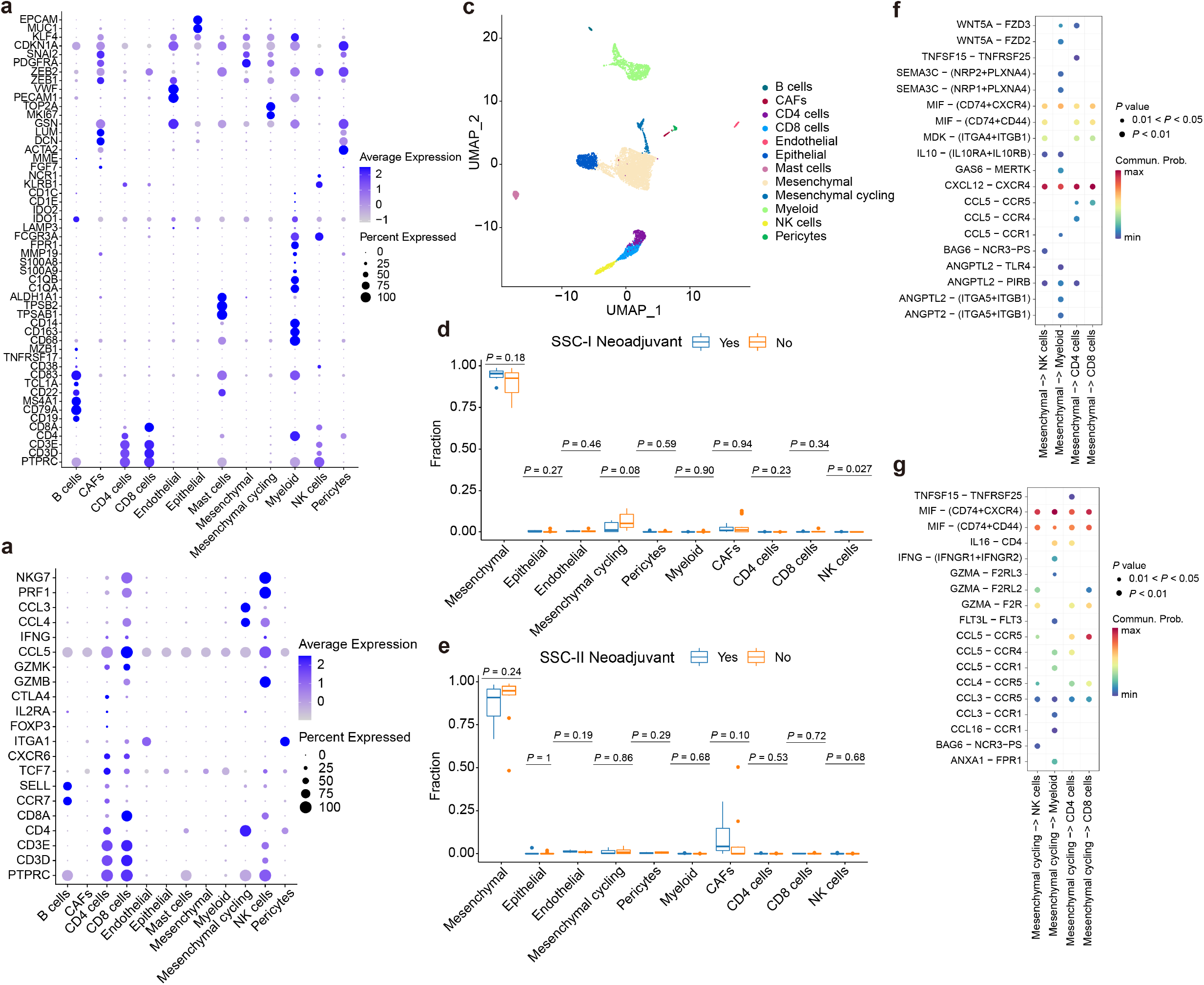
**(a-b).** Dot plots showing the **(a)** cell markers (except T cell markers) and **(b)** T cell markers expressions across the 12 cellular clusters. The size of dots represents the proportion of cells expressing the particular marker, and the spectrum of color indicates the mean expression levels of the markers. **(c).** UMAP plot of scRNA-seq profiles all SS patients, colored by cell types. **(d).** Cell type distributions in neoadjuvant and no neoadjuvant treated SSC-I patients after CIBERSORTx deconvolution. Middle line: median; box edges: 25th and 75th percentiles. Mann-Whitney U test. **(e).** Cell type distributions in neoadjuvant and no neoadjuvant treated SSC-II patients after CIBERSORTx deconvolution. Middle line: median; box edges: 25th and 75th percentiles. Mann-Whitney U test. **(f-g).** Ligand–receptor pairs between **(f)** mesenchymal and **(g)** mesenchymal cycling cells and immune cells (NK cells, myeloid cells, CD4 cells, and CD8 cells.

**Extended Data Figure. 14:**
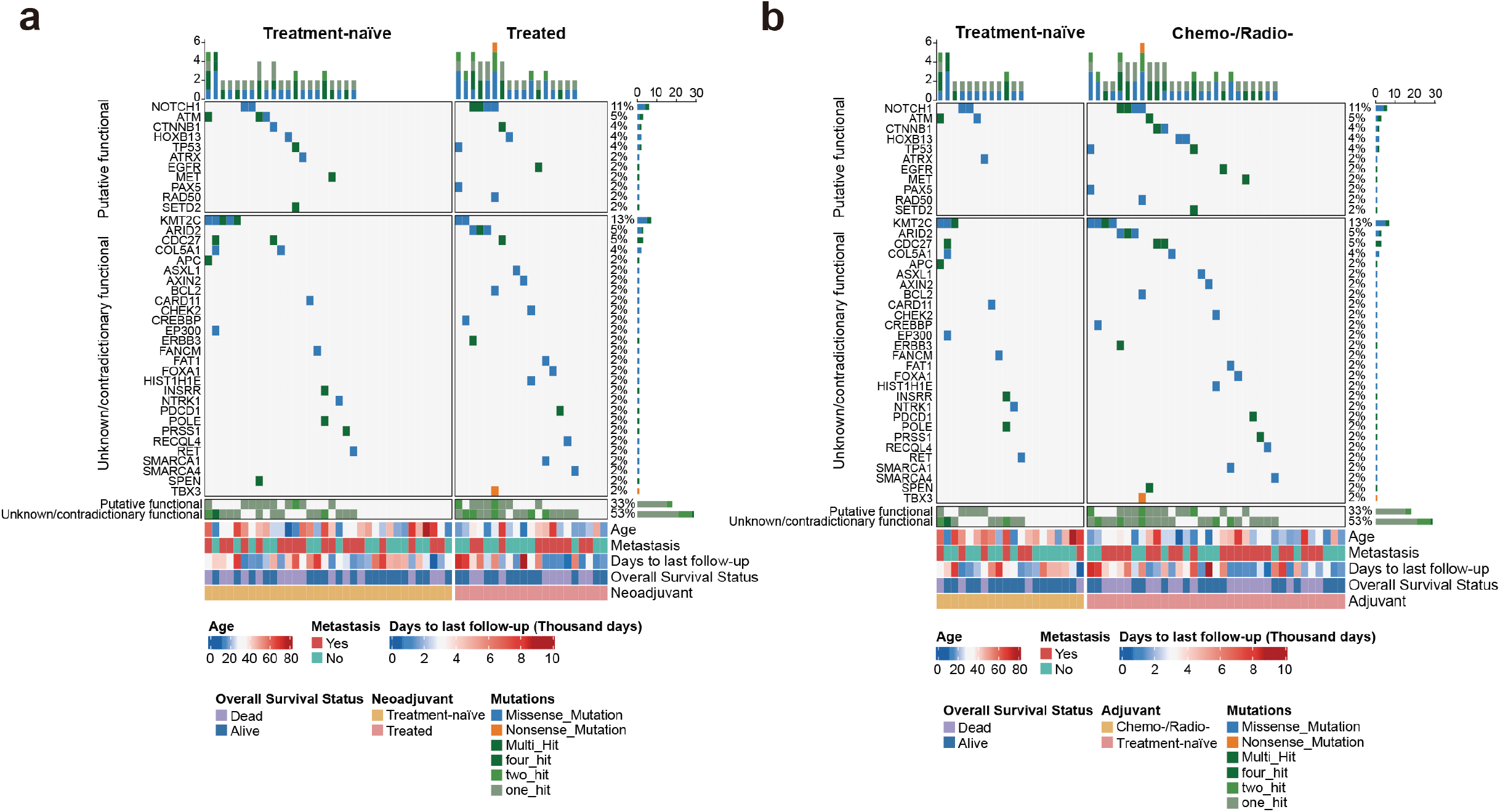
**(a-b).** Fifty-five synovial sarcoma patients with/without mutation data are ordered by their mutation frequencies and separated by **(a)** Neoadjuvant and **(d)** adjuvant treatment status.

**Extended Data Figure. 15:**
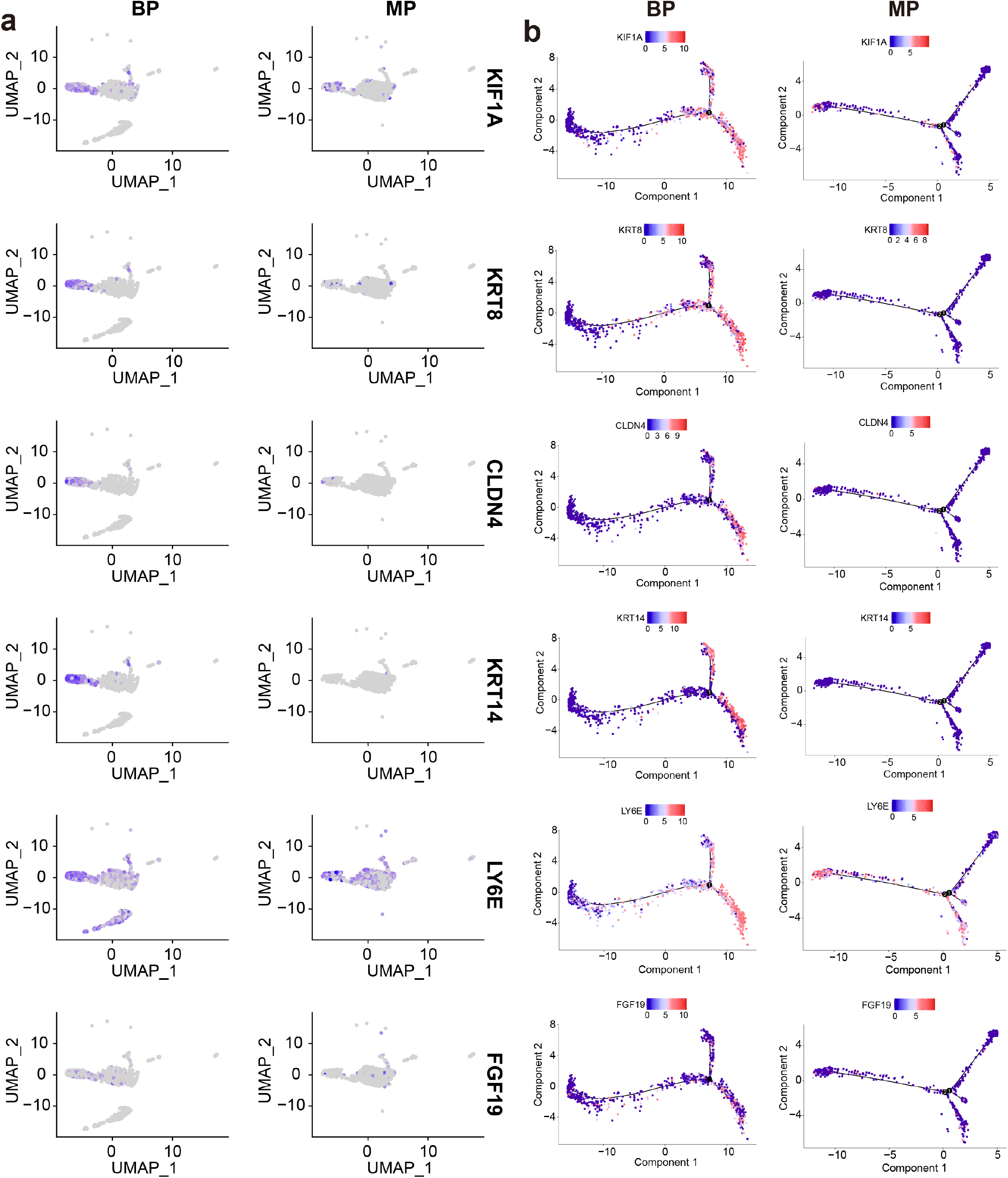
**(a).** Feature plots for MET key genes (KIF1A, KRT8, CLDN4, KRT14, LY6E and FGF19) The color legend shows the normalized expression levels of the genes. **(b).** The expression levels of the MET key genes (KIF1A, KRT8, CLDN4, KRT14, LY6E and FGF19) along with the pseudotime course of MET process as determined by the Monocle 2 trajectory analysis.

**Extended Data Figure. 16:**
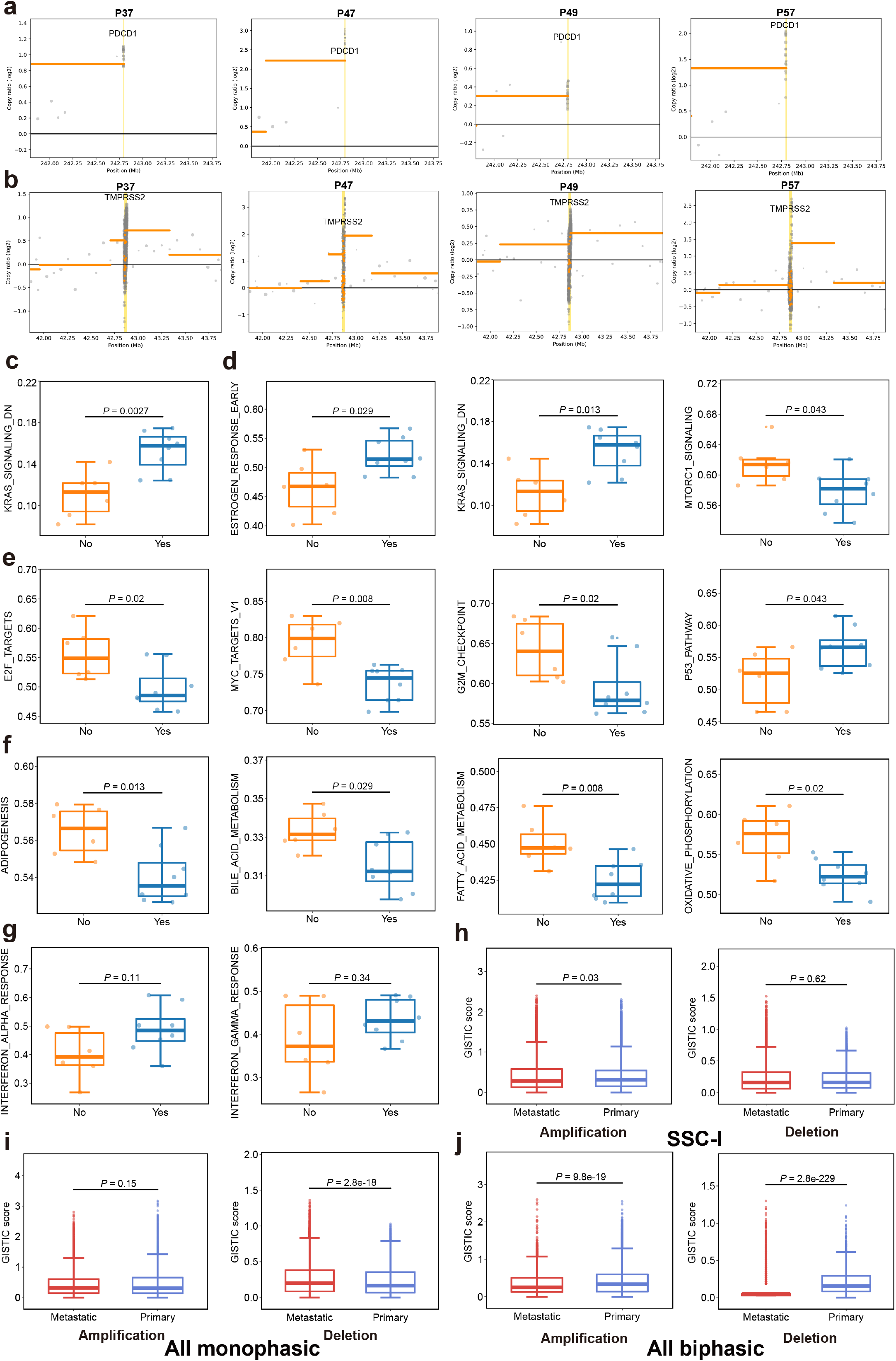
**(a-b).** Scatter plot of the copy ratio of **(a)** PD-1 and **(b)** TMPRSS2 in P37, P47, P49, and P57. **(c).** Distributions of the ssGSEA score of the KRAS signaling downregulated genes between biphasic patients with- and without-PD-1 high amplification. Middle line: median; box edges: 25th and 75th percentiles. Mann-Whitney U test. **(d-g).** Distributions of the ssGSEA score of the **(d)** signaling, **(e)** proliferative, **(f)** metabolic, and **(g)** immune-related gene sets between biphasic patients with- and without-TMPRSS2 high amplification. Middle line: median; box edges: 25th and 75th percentiles. Mann-Whitney U test. **(h-j).** GISTIC scores distributions of amplifications and deletions between primary and metastatic in **(i)** SSC-I, **(i)** all monophasic, and **(j)** all biphasic tumors. Middle line: median; box edges: 25th and 75th percentiles. Mann-Whitney U test.

## Supplementary table legends

**Supplementary table 1:** Clinical information of 55 patients in the discovery cohort.

**Supplementary table 2:** Identified fusion genes by FusionCatcher and STAR-fusion.

**Supplementary table 3:** PCR primers of the whitelist fusion genes.

**Supplementary table 4:** Cell marker for cell cluster annotations.

**Supplementary table 5:** List of well-known oncogenes in our gene panel (n = 389).

**Supplementary table 6:** Significantly enriched gene sets in the GSEA results.

## Notes

### Competing Interest Statement

The authors have declared no competing interest.

